# The evolutionarily conserved EHMT1/G9a histone methyltransferase family regulates sleep maintenance through ROS homeostasis in insulin-producing cells

**DOI:** 10.64898/2026.08.28.747754

**Authors:** Mireia Coll-Tané, Lara V. van Renssen, Nicholas Raun, Jie Han, Núria Ribas Ros, Boyd van Reijmersdal, Jenny Luong, Chiara Pignato, Franziska Kampshoff, Naihua N. Gong, Spencer G. Jones, Sigrid Pillen, Anna Castells-Nobau, Jordi Mayneris-Perxachs, Marieke Klein, Matthew S. Kayser, Tjitske Kleefstra, Annette Schenck

## Abstract

Sleep disturbances are a common, still poorly characterized feature of Kleefstra syndrome (KLEFS1), a neurodevelopmental disorder caused by rare variants in the epigenetic regulator *EHMT1*. The gap in understanding the characteristics and origin of these sleep disturbances poses a major barrier for therapy development. In this cross-species study, we reveal that 70% of individuals with KLEFS1 experience severe sleep maintenance insomnia, marked by fragmented sleep due to frequent night awakenings. Furthermore, common genetic variation at the *EHMT1* locus was associated with short sleep and insomnia symptoms in the general population. *Drosophila* mutants of the *EHMT1* orthologue *G9a* recapitulate these phenotypes, exhibiting reduced and fragmented sleep. We show that G9a is required in insulin-producing cells (IPCs) and the fat body, in the latter during development, to ensure adult sleep integrity. Untargeted metabolomics revealed widespread metabolic dysregulation in *G9a* mutants, particularly affecting methionine metabolism. Mutants exhibited reduced methionine and elevated methionine sulfoxide (Met-SO), pointing to increased reactive oxygen species (ROS). Redox sensors revealed increased H_2_O_2_-dependent oxidation in the larval brain and an elevated glutathione redox potential in IPCs during development but not in adulthood. IPC-specific knockdown of *MsrA*, the enzyme that reduces Met-SO back to methionine, reproduced sleep fragmentation. Developmental, but not acute, antioxidant treatment fully restored adult sleep consolidation, demonstrating that G9a safeguards sleep via ROS homeostasis in early life. Finally, we show that a *Drosophila* sleep-restriction paradigm based on human sleep-restriction therapy can override the developmental defects and restore sleep continuity in adulthood. Our findings establish an evolutionarily conserved role for EHMT1/G9a in sleep regulation and provide a mechanistic framework to understand and treat sleep disturbances in KLEFS1.

## Introduction

Sleep disturbances are among the most common co-occurring features of neurodevelopmental disorders (NDDs), affecting between 50 and 86% of individuals and substantially diminishing the quality of life of both affected individuals and their families^1^. Kleefstra syndrome (KLEFS1) is a monogenic NDD caused by heterozygous loss-of-function variants in the *euchromatic histone lysine methyltransferase 1 (EHMT1)* gene (OMIM #610253)^2,3^. The syndrome is characterized by moderate to severe intellectual disability, developmental delay, childhood hypotonia, and distinctive facial features^4–8^. Individuals with KLEFS1 also have a high prevalence of comorbidities, including autism spectrum disorder, overweight or obesity, and behavioral problems^4,7^. Sleep problems in KLEFS1 have recently gained increasing attention, triggered by the observation that phases of severe sleep problems may precede regressive episodes^4,9^. Our first studies registering sleep as part of broader clinical and neurodevelopmental phenotyping reported incidences of sleep problems in 46-79% of KLEFS1 individuals and documented that these can substantially impact health and daily functioning^4,7,9,10^. Insomnia appears to be the most prevalent sleep disturbance^2^, but so far no dedicated standardized sleep assessments of cohorts have been reported. Moreover, despite their emerging prevalence and clinical importance, the biological underpinnings of sleep disturbances in KLEFS1 remain unknown.

*EHMT1* encodes an evolutionarily conserved lysine methyltransferase that regulates chromatin state and gene expression^3,6,11^. In *Drosophila melanogaster*, the fruit fly, G9a is the common ancestral ortholog of the mammalian EHMT1 and EHMT2 proteins, which co-operate in a protein complex. The protein family exerts established roles in neurodevelopment, memory, and adaptation to environmental stress^11–14^. The evolutionary conservation of EHMT1/G9a, together with the established contribution of *Drosophila* to the study of both neurodevelopmental disorders and sleep physiology^15–18^, provides an opportunity to dissect the role of this epigenetic regulator in sleep regulation.

In this cross-species and cross-disciplinary study, we integrated clinical sleep assessments and human genetic association analysis with genetics, behavioral, metabolomics and imaging analyses in *Drosophila* to investigate the contribution of *EHMT1/G9a* to sleep regulation. We first characterized sleep disturbances in individuals with KLEFS1 and examined whether common genetic variation at the *EHMT1* locus is associated with sleep-related traits in the general population. We then used *Drosophila* to identify the tissues and developmental period through which G9a influences sleep, and to investigate the molecular processes and mechanisms underlying this phenotype. Finally, building on the obtained insights, we tested and identified pharmacological and behavioral approaches to improve sleep in our preclinical model, with the latter holding significant translational potential.

## Results

### Individuals with KLEFS1 show high rates of night waking

To obtain insights into the prevalence and nature of sleep disturbances in KLEFS1, we collected qualitative and/or quantitative sleep data from 33 individuals. Parents or caregivers completed an extensive sleep behavior questionnaire (the Modified Simonds & Parraga Sleep Questionnaire)^19–21^. Cognitive impairment was the most prevalent comorbidity in this cohort (**Supplementary Fig. 1a**). 70% (23 out of 33) of parents/caregivers reported that their child was experiencing sleep disturbances at the time of completing the questionnaire. These disturbances primarily involved nighttime awakenings (19/23; 82.6%) and, to a lesser extent, early awakenings (10/23; 43.5%) (**Fig. 1a**). Notably, none of the participants reported difficulty falling asleep.

**Figure 1.**
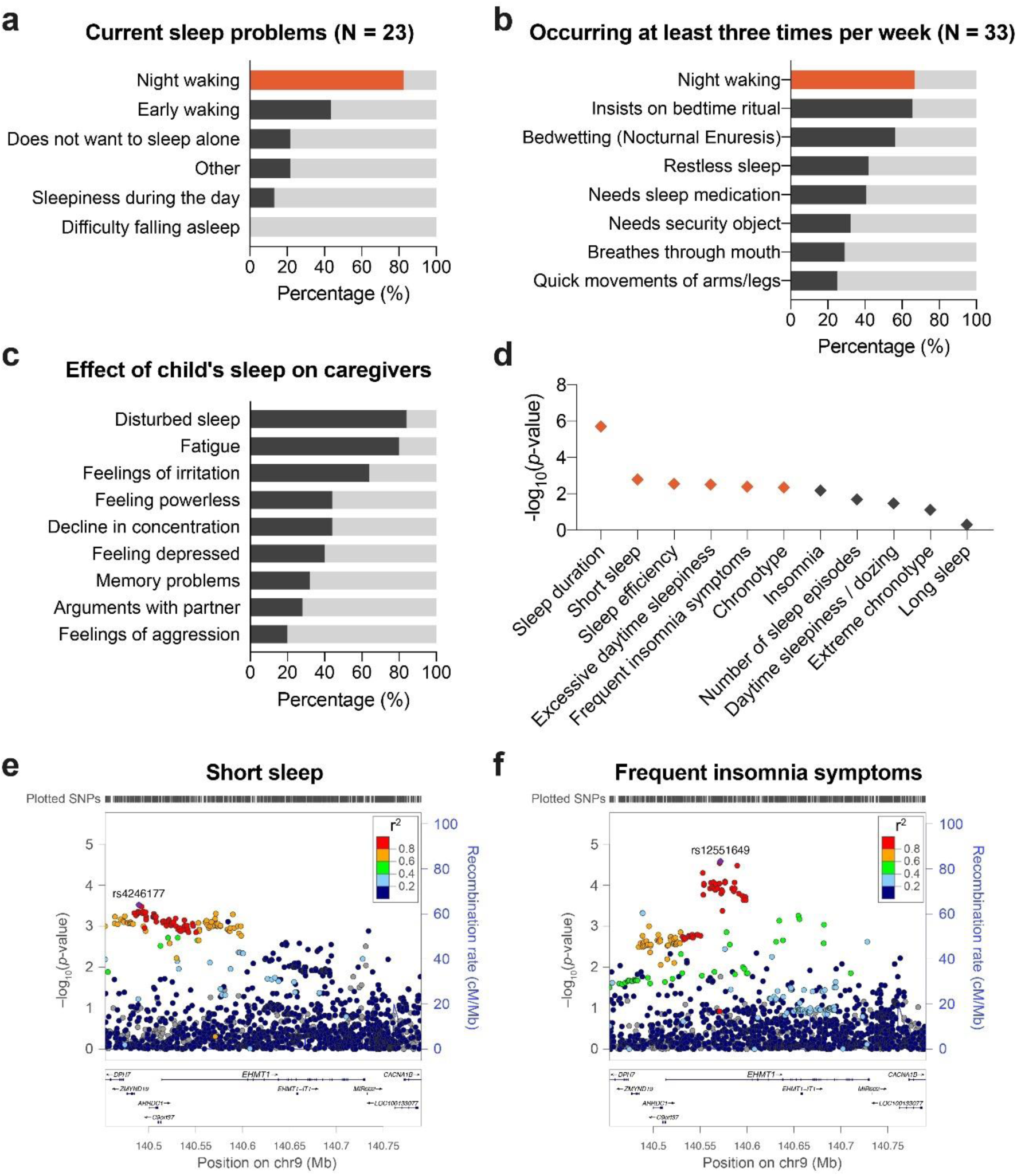
Rare and common variants in *EHMT1* determine sleep integrity and duration. **(a)** Nature of the sleep problems reported by the parents of individuals with KLEFS1 who indicated their child to currently have disturbed sleep (23 out of 33 individuals). **(b)** Sleep disturbances reported to occur at least once a week in the whole cohort (n = 33), independently of whether the parents consider their child to currently experience sleep problems. (**a,b**) The colored bar highlights night wakening as the most disturbed aspect of sleep in KLEFS1 **(c)** Effect of child’s sleep on family members (n = 25). **(d)** Common variants in *EHMT1* are significantly associated with multiple sleep traits. Colored diamonds withstand multiple testing correction (*p* < 0.00455). **(e)** Regional association plots showing association signals for “short sleep” and **(f)** “frequent insomnia symptoms” at the *EHMT1* locus. Data are shown as −log_10_(*p*-value) for individual SNPs. The color of each marker reflects its linkage disequilibrium (r^2^) with the strongest associated SNP indicated as a purple diamond. The recombination rate is indicated in blue. Chr, chromosome; cM, centimorgan; Mb, megabase.

As the International Classification of Sleep Disorders 3 (ICSD-3) criteria defines a sleep disorder as symptoms occurring at least three times per week^19,22^, we applied this criterion regardless of whether caregivers reported current sleep problems. The most frequently reported issue was nighttime awakenings occurring three or more times per week (22/33; 67%) (**Fig. 1b**), consistent with a sleep disorder characterized by sleep-maintenance insomnia. On average, most individuals with KLEFS1 woke twice per night (12/33; 36.4%), whereas others woke once (11/33; 33.3%) or three times (9/33; 27.3%) per night (**Supplementary Fig. 1b**). The majority of the KLEFS1 cohort fell back asleep within a few (16/26; 61.5%) or up to 30 minutes (11/26; 42.3%) (**Supplementary Fig. 1c**).

In addition to the questionnaires, caregivers were asked to provide an annotated graphical sleep diary^19–21^ for two consecutive weeks. We included only 16 diaries in the final analysis, selecting those that were complete and for which parents indicated reliable awareness of their child’s nighttime awakenings. Analysis of these diaries corroborated the questionnaire findings. Nine of 16 KLEFS1 individuals exhibited increased wake after sleep onset (WASO), particularly the youngest participants (3-4 years) and post-pubescent individuals (≥18 years) (**Table 1**). We also observed an increased number of nighttime awakenings in 8 of 16 participants, with a similar age distribution to that observed for WASO (**Table 1**). These findings indicate that increased WASO reflects multiple awakenings rather than a single prolonged wake episode, consistent with sleep fragmentation, in agreement with the questionnaire data (**Supplementary Fig. 1b**). We also observed reduced total sleep time in 3 of 16 individuals (**Table 1**); however, in 2 of these individuals, this reduction was, in terms of sleep duration, compensated for by daytime naps (**Table 1**). Additionally, 4 of 16 individuals showed a modest increase in sleep onset latency (33-41 minutes).

**Table 1.** Quantitative data from the sleep diaries of individuals with KLEFS1. Quantitative sleep parameters from parent-reported sleep-wake diaries confirmed the presence of increased WASO and increased number of night awakenings. Some individuals with KLEFS1 also showed decreased total sleep time and increased SOL. Values in bold deviate from established healthy reference values (see Materials & Methods).

| Individual | 1 | 2 | 3 | 4 | 5 | 6 | 7 | 8 | 9 | 10 | 11 | 12 | 13 | 14 | 15 | 16 |
| --- | --- | --- | --- | --- | --- | --- | --- | --- | --- | --- | --- | --- | --- | --- | --- | --- |
| Sex | F | M | F | F | M | F | F | F | M | F | F | M | F | M | F | F |
| Age (years) | 3 | 3 | 3 | 4 | 4 | 5 | 6 | 7 | 11 | 16 | 18 | 20 | 23 | 26 | 26 | 33 |
| **Sleep wake calendar (averages)** |  |  |  |  |  |  |  |  |  |  |  |  |  |  |  |  |
| Days measured | 14 | 12 | 14 | 7 | 14 | 14 | 14 | 14 | 27 | 14 | 13 | 14 | 16 | 12 | 12 | 14 |
| Time in bed (hh:mm) | 10:39 | 11:23 | 11:12 | 11:04 | 11:16 | 11:35 | 10:55 | 9:57 | 10:21 | 8:17 | 9:20 | 10:16 | 10:37 | 11:27 | 11:48 | 10:09 |
| Total Sleep Time (Night) (hh:mm) | **9:46**<br>(↓) | 10:56 | **9:07**<br>(↓) | 10:12 | 10:00 | 10:53 | 10:40 | 9:40 | 10:07 | **7:51**<br>(↓) | 8:11 | 8:53 | 7:30 | 9:46 | 10:57 | 8:55 |
| Total Sleep Time (including naps) (hh:mm) | 10:32 | - | **9:22**<br>(↓) | - | - | - | - | - | - | - | - | - | 7:56 | - | - | - |
| Sleep Onset Latency (min) | 21 | 5 | **33**<br>(↑) | **34**<br>(↑) | 2 | 21 | 9 | 7 | 8 | 18 | 0 | 9 | **39**<br>(↑) | 18 | 0 | **41**<br>(↑) |
| Wake After Sleep Onset (WASO; min) | 19 | 20 | **92**<br>(↑) | 12 | **50**<br>(↑) | 20 | 4 | 6 | 3 | 1 | **69**<br>(↑) | **69**<br>(↑) | **106**<br>(↑) | **43**<br>(↑) | **37**<br>(↑) | 12 |
| Wake After Sleep Offset (WASF; min) | 12 | 2 | 0 | 4 | 23 | 0 | 0 | 3 | 1 | 6 | 0 | 4 | **40**<br>(↑) | **38**<br>(↑) | 13 | 19 |
| WASO + WASF (min) | **32**<br>(↑) | 22 | **92**<br>(↑) | 17 | **73**<br>(↑) | 20 | 4 | 10 | 5 | 7 | **69**<br>(↑) | **73**<br>(↑) | **147**<br>(↑) | **82**<br>(↑) | **51**<br>(↑) | **32**<br>(↑) |
| Number of night awakenings | **1.1**<br>(↑) | 0.9 | **3.1**<br>(↑) | 0.9 | **2.9**<br>(↑) | 0.9 | 0.2 | **1.4**<br>(↑) | 0.1 | 0.1 | **1.3**<br>(↑) | **4.0**<br>(↑) | **2.8**<br>(↑) | 0.7 | **2.5**<br>(↑) | 0.4 |
| Sleep Efficiency (SE; %) | 91.6 | 96.0 | **81.4**<br>(↓) | 92.2 | 88.8 | 93.9 | 97.9 | 97.0 | 97.9 | 94.8 | 87.9 | 86.5 | **70.3**<br>(↓) | 85.2 | 92.8 | 87.9 |

Finally, the sleep questionnaires also allowed us to evaluate the impact of the child’s sleep on their caregivers. 84% of responding caregivers (21/25) reported that their sleep was disturbed by their child, with 80% experiencing fatigue due to the child’s sleep problems (20/25), and approximately 65% reporting irritability (16/25) (**Fig. 1c**). More than 40% of caregivers reported feelings of powerlessness (11/25) and/or depression (10/25), as well as reduced concentration (11/25). Approximately one-third of caregivers reported memory problems (8/25), increased arguments with their partners (7/25), and/or feelings of aggression (5/25) (**Fig. 1c**). Together, these findings highlight the high prevalence of sleep disturbances, particularly sleep fragmentation, in individuals with KLEFS1, as well as their substantial negative impact on family well-being.

### Common variants in *EHMT1* are associated with multiple sleep traits

We next asked whether genetic variation of *EHMT1* likewise impacts sleep properties in the general population. We therefore determined whether common single nucleotide polymorphisms (SNPs) in *EHMT1,* detected in multiple genome-wide association studies (GWAS), show significant associations with sleep. We retrieved gene-based *p*-values for *EHMT1* (NCBI Gene ID: 79813) for all available GWAS in the GWAS atlas^23^. After correction for the number of sleep traits examined, *EHMT1* showed significant associations with “sleep duration”, “short sleep”, “sleep efficiency”, “excessive daytime sleepiness”, “frequent insomnia symptoms”, and “chronotype” (**Fig. 1d-f**, **Supplementary Fig. 2a-d**, and **Supplementary Table 1**). In conclusion, SNPs in *EHMT1* are associated with sleep traits closely related to the sleep disturbances found in individuals with KLEFS1, supporting that EHMT1 regulates sleep in health and disease.

### The EHMT1 ortholog, G9a, is required for sleep integrity in *Drosophila*

To establish an animal model that allows dissecting the molecular and systemic mechanisms underlying sleep regulation by *EHMT1*, we assessed sleep in *Drosophila* carrying a null allele of the *EHMT1* ortholog *G9a*^11^. Hemizygous *G9a* null mutant males showed a significant decrease in total sleep time both during the day (light period, LP) and night (dark period, DP) compared with isogenic controls. Because also in *Drosophila*, not only total sleep time but also sleep consolidation, particularly of nighttime sleep episodes are important for brain physiology and function, including learning^24–26^, we analyzed sleep architecture.

This revealed a dramatically shorter mean sleep-bout duration during both day (to 53%, average of 30 min in controls versus 16 min in *G9a* mutants) and night (to 42%, average of 200 min versus 83 min) (**Fig. 2a,b**). In contrast, *G9a* mutants exhibited a significantly increased number of sleep bouts (**Fig. 2c**), which however only partly compensated for the reduced bout duration (**Fig. 2a**). We also assessed homozygous *G9a* null mutant females. These showed decreased and fragmented sleep to an extent similar to the hemizygous mutant males (**Fig. 2a’-c’**). By contrast, females carrying only one *G9a* null allele did not show significant differences in sleep quantity or quality (**Fig. 2a’-c’**), indicating that G9a function needs to fall short of a critical threshold for sleep defects to manifest.

**Figure 2.**
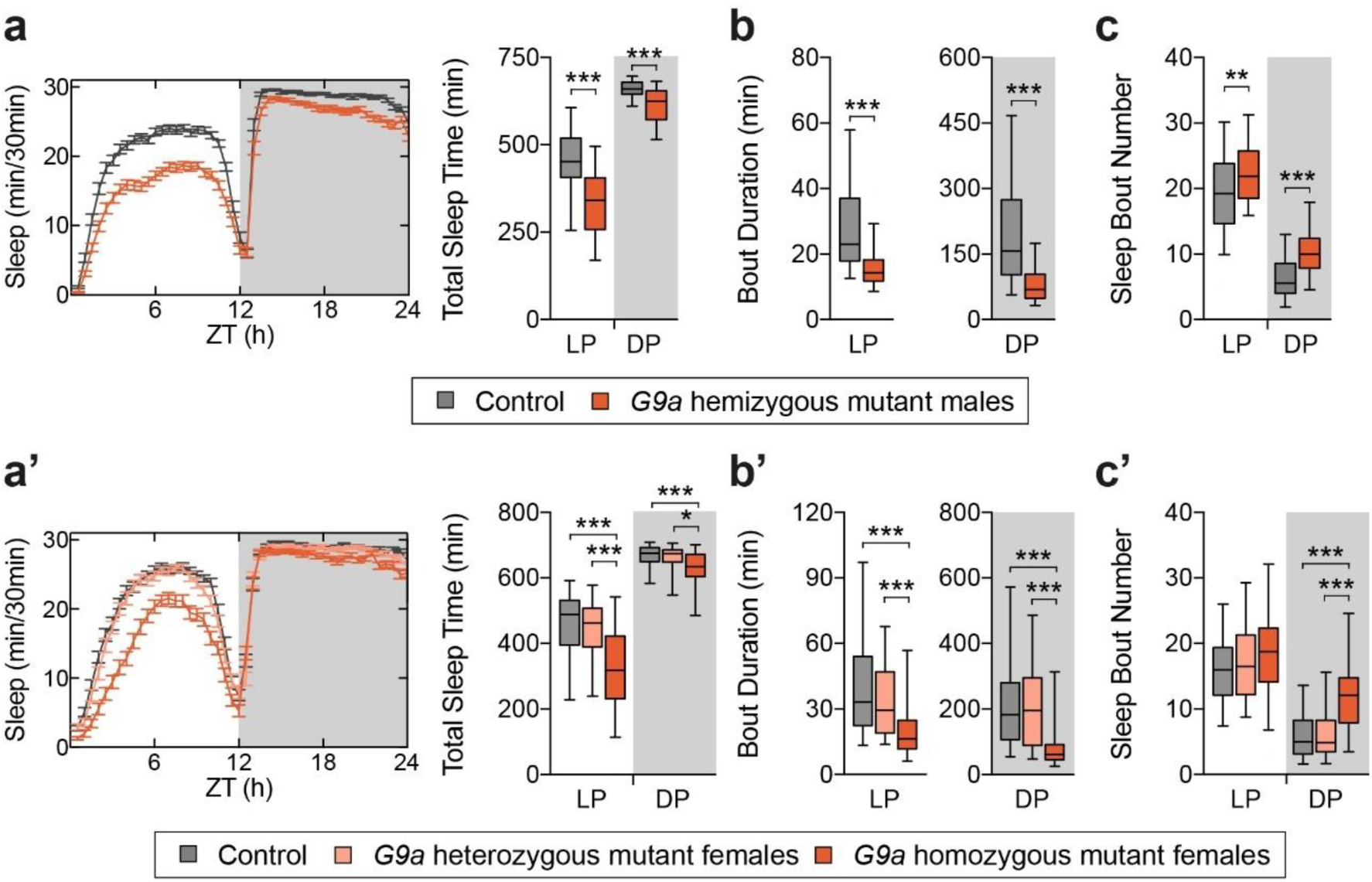
Loss of *G9a* leads to decreased and fragmented sleep, and an increased latency to fall back to sleep when wakened. **(a,a’)** Sleep profiles and quantification. **(b,b’)** Average duration and **(c,c’)** number of sleep bouts in the light period (LP, ZT0-12) and dark period (DP, ZT12-24). (a) *G9a* hemizygous mutant male flies (n = 92) show decreased total sleep time during both the LP and DP (*p* < 0.0001) compared to isogenic controls (n = 90). **(b)** Average bout duration in *G9a* hemizygous mutant male flies is strongly reduced both in the LP and DP (*p* < 0.0001), whilst **(c)** the number of bouts is increased (*p* = 0.0012 and *p* < 0.0001). Two-tailed unpaired Welch’s t-test. Bonferroni correction for multiple testing. (a’-c’) *G9a* homozygous mutant female flies (n = 58) show decreased sleep time in the LP and DP compared to isogenic controls (n = 69; *p* < 0.0001 and *p* = 0.0006) and *G9a* heterozygous mutant female flies (n = 53; *p* < 0.0001 and *p* = 0.01). *G9a* homozygous mutant female flies show reduced bout length during the LP and DP (*p* < 0.0001), accompanied by increased bout number during the DP (*p* < 0.0001). Data are presented as boxplots (25^th^-75^th^ percentiles, median; whiskers indicate 5^th^-95^th^ percentiles). Kruskal-Wallis test with Dunn’s correction. Bonferroni correction for multiple testing. *p*-values are indicated as follows: \**p* ≤ 0.05, \*\**p* ≤ 0.01, \*\*\**p* ≤ 0.001.

We next asked whether G9a loss disturbs the sleep homeostat. To assess this, we measured sleep rebound following overnight mechanical sleep deprivation. Both hemizygous *G9a* mutants and their controls showed a robust daytime sleep rebound (**Supplementary Fig. 3a,b**), indicating that the homeostatic response to sleep deprivation was preserved in mutants. We also addressed whether G9a regulates circadian rhythm by assessing the amplitude of rest:activity rhythms in constant darkness using fast Fourier transform analysis^27^. We found that *G9a* mutants exhibited circadian rhythmicity similar to controls in free-running conditions (**Supplementary Fig. 3c,c’**). Together, our data reveal that *Drosophila* G9a, like its human orthologue EHMT1, regulates sleep quantity and integrity independently of sleep homeostasis and circadian rhythmicity.

### G9a is required in insulin-producing cells and fat body for sleep integrity

We next sought independent support for the role of G9a in sleep and aimed to identify the tissues or cells through which it exerts its regulatory function. Using the UAS-Gal4 system and inducible RNAi^27,28^, we knocked down *G9a* in specific cell-types using a previously molecularly and phenotypically validated *UAS-G9a^RNAi^*line^6,13^. Because G9a is required in the mushroom body (MB) for short- and long-term memory formation^29^, and this structure also plays a critical role in sleep regulation^29,30^, we first targeted *G9a* knockdown to the MB. However, MB-specific *G9a* knockdown using the *R14H06*-Gal4 driver^31^ did not alter sleep quantity or architecture relative to both parental controls (**Supplementary Fig. 4a-a’’**).

We previously reported that, upon oxidative stress (OS) exposure, G9a limits the cost-intensive transcriptional activation of OS-resistance genes thereby preserving energy stores and metabolic homeostasis^32^. As both OS and energy metabolism have previously been linked to sleep regulation^33^, we asked whether G9a regulates sleep in the same cells in which it controls energy homeostasis, namely the insulin-producing cells (IPCs) and the fat body (FB)^33–35^. In *Drosophila*, IPCs are crucial regulators of metabolic and sleep homeostasis^33–35^, whereas the FB, the analog of the mammalian liver and adipose tissue, is a key organ for energy storage and homeostasis and for immune function^36,37^. *G9a* knockdown in IPCs using the *dilp2-Gal4* driver recapitulated the strongly fragmented nighttime sleep observed in *G9a* null mutants, characterized by a decrease in sleep bout duration and a higher number of sleep bouts compared to both parental controls (**Fig. 3a-a’’**). Additionally, knockdown of *G9a* in the FB with the *C7-Gal4* driver also led to nighttime sleep fragmentation (**Fig. 3b-b’’**). Total sleep amount relative to both parental controls was not significantly changed upon knockdown of *G9a* in either tissue (**Fig. 3a,b**). We validated these findings using independent IPC- and FB-specific drivers, *R96A08-Gal4* and *FB-Gal4,* respectively (**Supplementary Fig. 4b-c’’**). In conclusion, G9a regulates sleep architecture in cells also critical for metabolic homeostasis regulation.

**Figure 3.**
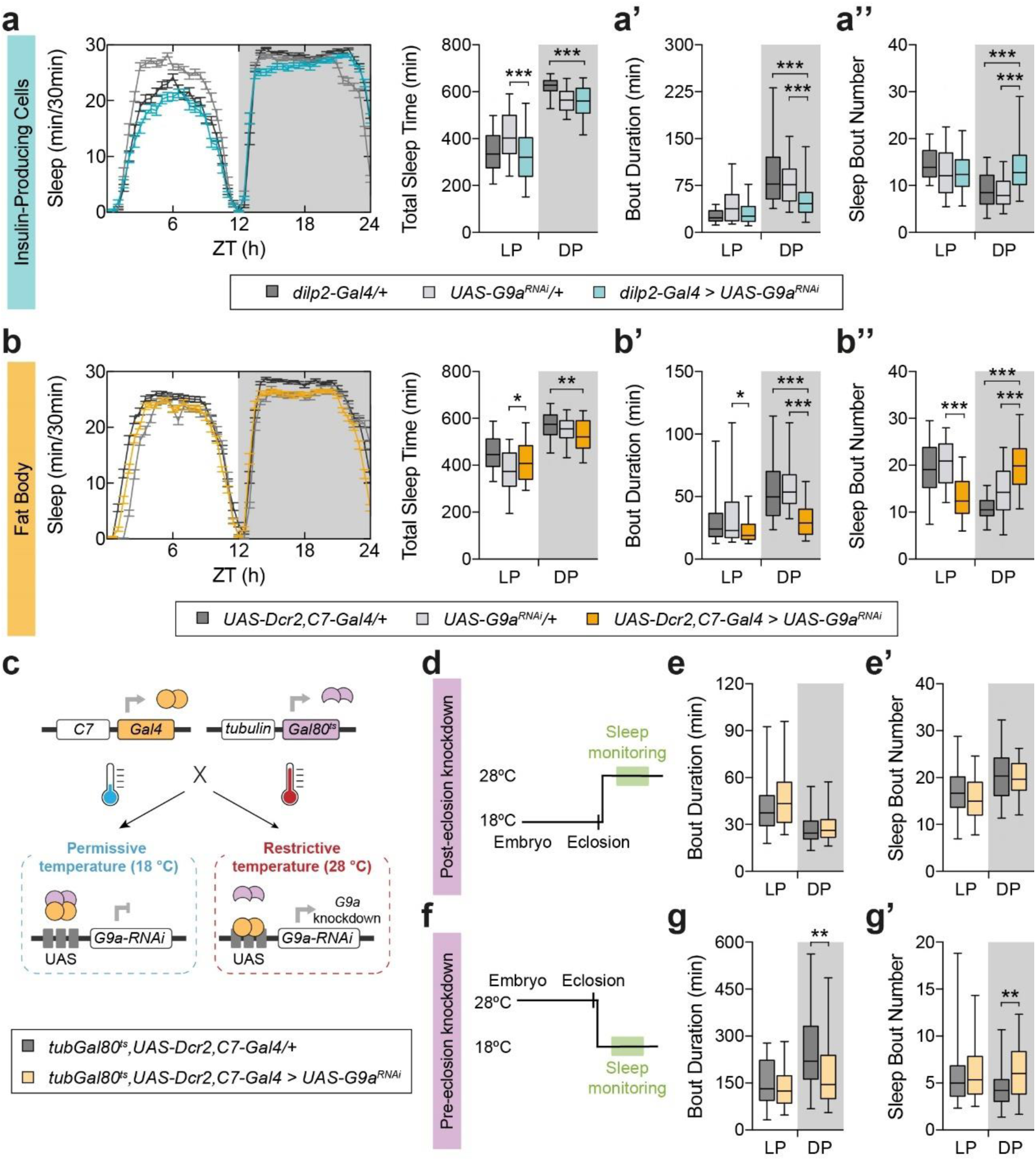
G9a is required in insulin-producing cells and fat body for adult sleep integrity, in the latter during development. **(a-e)** Sleep profiles and quantification. **(a’-e’)** Average duration and **(a’-e’’)** number of their sleep bouts in the light period (LP, ZT0-12) and dark period (DP, ZT12-24). (a-a’’) *G9a* knockdown in insulin-producing cells (*dilp2-Gal4 > UAS*-*G9a^RNAi^*, n = 72) leads to shorter sleep bouts (*p* < 0.0001) accompanied by an increase in their number (*p* < 0.0001) during the DP when compared to parental controls (*dilp2-Gal4/+*, n = 72; *UAS*-*G9a^RNAi^/+*, n = 72). (b-b’’) *G9a* knockdown in the fat body (*UAS-Dcr2,C7-Gal4 > UAS*-*G9a^RNAi^*, n = 74) leads to shorter sleep bouts (*p* < 0.0001) accompanied by an increase in their number (*p* < 0.0001) during the DP when compared to parental controls (*UAS-Dcr2,C7-Gal4/+* controls in dark grey, n = 74; *UAS*-*G9a^RNAi^/+* controls in light grey, n = 84). Kruskal-Wallis test with Dunn’s correction. Bonferroni correction for multiple testing. **(c)** Schematic representation of the TARGET system. **(d)** Temperature shifts used to induce post-eclosion *G9a* knockdown, restricting it to adulthood. **(e)** Average sleep bout duration and **(e’)** number of sleep bouts in flies with *G9a* knockdown in the fat body exclusively during post-eclosion stages (*tub-Gal80^ts^, C7-Gal4, UAS-Dcr2 > UAS*-*G9a^RNAi^*, n = 103) compared to isogenic controls (*tub-Gal80^ts^, C7-Gal4, UAS-Dcr2/+*, n = 73). Post-eclosion *G9a* knockdown does not affect sleep bout architecture. **(f)** Temperature shifts used to induce pre-eclosion *G9a* knockdown, restricting it to adulthood. **(g)** Average sleep bout duration and **(g’)** number of sleep bouts in flies with *G9a* knockdown in the fat body exclusively during pre-eclosion stages (*tub-Gal80^ts^, C7-Gal4, UAS-Dcr2 > UAS*-*G9a^RNAi^*, n = 49) compared to isogenic controls (*tub-Gal80^ts^, C7-Gal4, UAS-Dcr2/+*, n = 50). *G9a* knockdown during developmental stages prior to eclosion is necessary and sufficient to cause fragmented sleep. Data are presented as boxplots (25^th^-75^th^ percentiles, median; whiskers indicate 5^th^-95^th^ percentiles). *p*-values are indicated as follows: \**p* ≤ 0.05, \*\**p* ≤ 0.01, \*\*\**p* ≤ 0.001.

### G9a is required in the developing fat body for adult sleep integrity

To determine whether G9a is required for sleep integrity early in development or acutely regulates sleep in adulthood, we took advantage of the temporal and regional gene expression targeting (TARGET) system^38^ to temporally restrict *G9a* knockdown (**Fig. 3c**). Flies carrying the *C7-Gal4*, UAS-G9a^RNAi^ and the ubiquitous, temperature-sensitive Gal4 repressor *tub-Gal80^ts^* were raised at a restrictive (18°C, to suppress *G9a* knockdown) or permissive temperature (28°C, facilitating knockdown). Flies maintained at 18°C throughout their lifespan showed comparable sleep architecture to their genetic background controls (**Supplementary Fig. 5a-b’**). Restricting *G9a* knockdown only to adulthood (shifting flies from 18 to 28°C upon eclosion) did not affect sleep quantity or architecture (**Fig. 3d-e’**). By contrast, restricting *G9a* knockdown to developmental stages (28°C pre-eclosion) resulted in decreased and fragmented nighttime sleep in the adult (**Fig. 3f-g’**), phenocopying the defects of *G9a* knockdown throughout the lifespan. Together, these results reveal that G9a activity in the developing FB contributes to the establishment of normal sleep consolidation in adulthood.

### G9a loss causes broad metabolic dysregulation and highly increased methionine oxidation

Having mapped the origin of sleep regulation to cells/tissues essential for systemic metabolic homeostasis, we performed unbiased untargeted metabolomics on *G9a* mutant fly heads to identify candidate metabolites mediating these effects (**Fig. 4a** and **Supplementary Table 2**).

**Figure 4.**
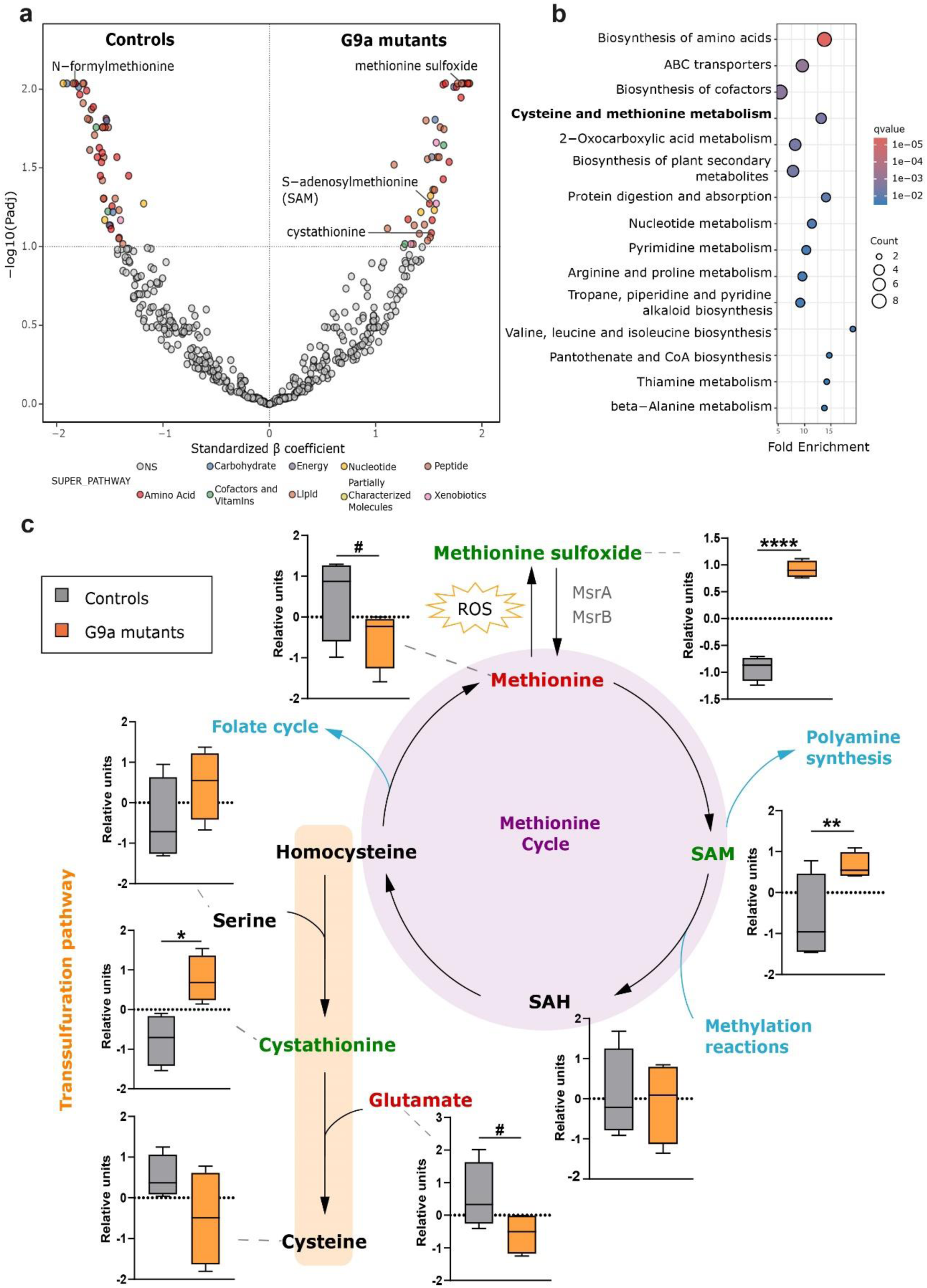
G9a loss leads to broad metabolic deregulation and highly increased methionine oxidation. **(a)** Volcano plot of differential metabolites found between the G9a mutant and control group identified employing robust linear regression models. Standardized β coefficient and log10 P values adjusted for multiple comparisons are plotted. **(b)** Dot-plot of significant (q-values < 0.1) over-represented KEGG pathways identified from significantly differential metabolites between the G9a mutant and control groups. The 15 more significant pathways are plotted. **(c)** Schematic representation of Met utilization pathways and metabolites from these pathways detected in our metabolomics analysis. Abbreviations: reduced Reactive Oxygen Species (ROS); S-adenosylmethionine (SAM); S-adenosylhomocysteine (SAH). Data from metabolites intensities normalized, transformed and scaled are represented as boxplots that extend from the 25^th^ to the 75^th^ percentiles with the median indicated. Whiskers indicate min and max values. Two-tailed unpaired Welch’s t-test. *p*–values are indicated as follows: #p≤ 0.1, \**p* < 0.05, \*\**p* < 0.01, \*\*\**p* < 0.001, \*\*\*\**p* < 0.0001.

Examination of the metabolomics data revealed that the methionine and cysteine metabolic pathways were strongly perturbed (**Fig. 4b** and **Supplementary Table 3**). Among 95 dysregulated metabolites, one of the most significantly increased was methionine sulfoxide (**Fig. 4a**) (methionine-SO), the primary oxidation product of methionine by ROS^39^. Methionine-SO levels were markedly elevated in G9a mutants, as indicated by a regression coefficient (β) of 1.82, whereas methionine levels were decreased (β = −1.17). This is noteworthy because methionine and cysteine are among the amino acids most susceptible to ROS-mediated oxidation^39,40^ and play central roles in maintaining cellular redox homeostasis^41^.

The essential amino acid methionine is additionally required to synthesize cysteine by trans-sulfuration^41^. Cysteine levels are reduced in *G9a* mutants (β = −1); however, this change did not reach statistical significance (**Fig. 4c**). This pathway also supports the biosynthesis of multiple antioxidants, such as glutathione and taurine^41,42^, raising the possibility that redox homeostasis is partly or more widely perturbed in *G9a* mutant conditions.

### *G9a* mutants show increased H_2_O_2_-dependent oxidation in the brain and altered glutathione redox potential in insulin-producing cells during development

Given the metabolic changes observed in *G9a* mutants, we next assessed cellular redox status using genetically encoded biosensors. These measurements are based on the expression of a redox-sensitive GFP (roGFP) that displays excitation-dependent fluorescence according to its oxidation state. Fusion of roGFP to the peroxidase Orp1 produces a hydrogen peroxide (H₂O₂) sensitive probe, whereas fusion to glutaredoxin (Grx1) allows changes in glutathione redox potential to be measured^43^. We selected roGFP-Orp1 to test whether the increased methionine sulfoxide detected by the metabolomics was accompanied by increased H_2_O_2_-dependent oxidation. Imaging of cytosolic roGFP2-Orp1 in larval *G9a* mutant brains indeed revealed greater oxidation (**Fig. 5a,a’**). Oxidation in the FBs of the same animals was unaffected (**Fig. 5b,b’**), revealing a tissue-specific role of G9a in H_2_O_2_-dependent oxidation.

**Figure 5.**
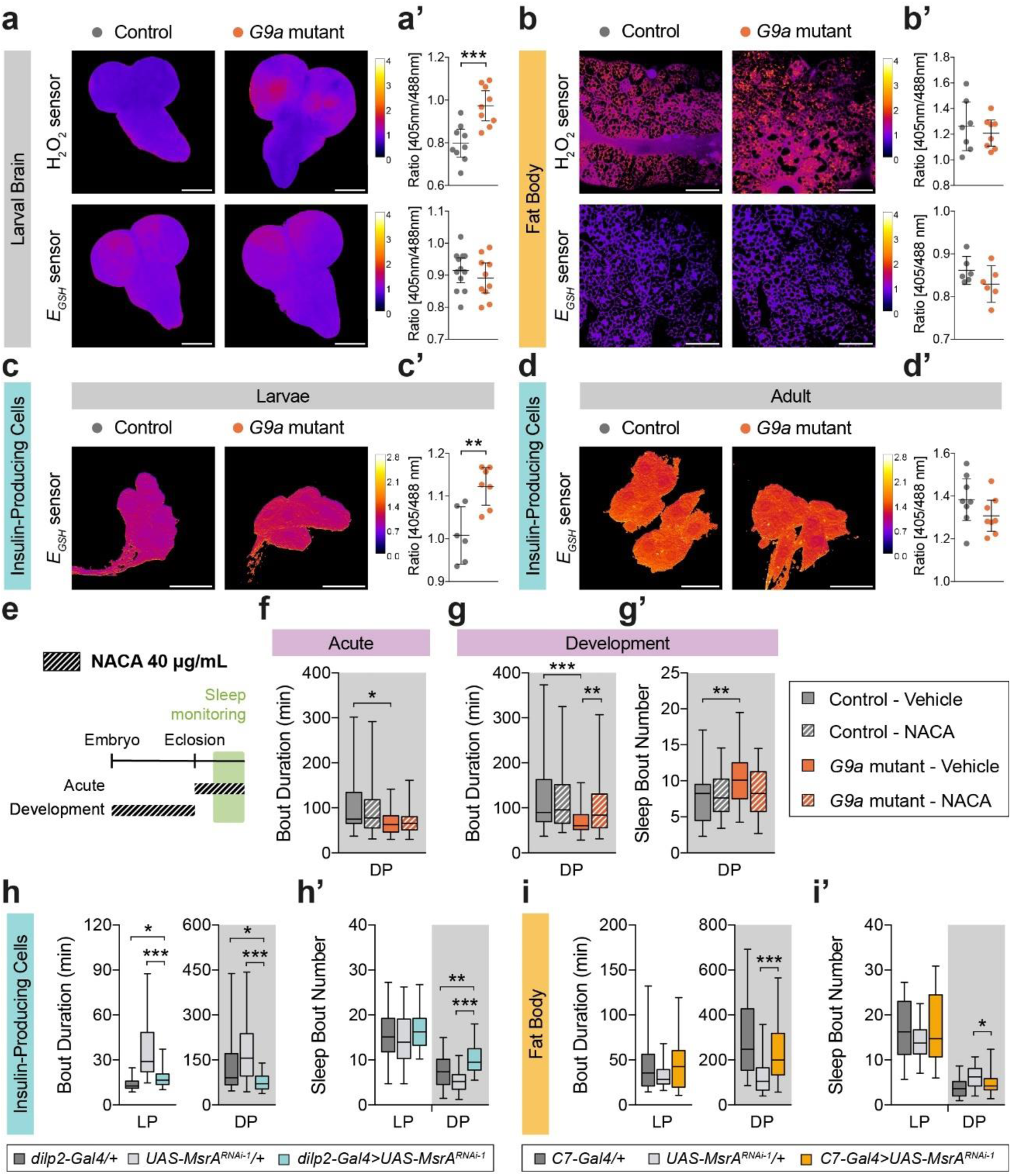
Increased ROS, in insulin-producing cells during development, underlies the nighttime sleep fragmentation in adult G9a mutants. **(a,a’)** Representative ratio images of whole larvae brains (*G9a* mutants and isogenic controls) expressing either cyto-roGFP2-Orp1 (H_2_O_2_ sensor) or cyto-Grx1-roGFP2 (*E_GSH_* sensor), and corresponding quantification. *G9a* mutants exhibit significantly elevated brain H_2_O_2_ levels (*p* = 0.0007). Scale bars: 150 μm. 9-12 flies per genotype. **(b,b’)** Representative ratio images of larval fat bodies expressing the same probes, and corresponding quantification. Scale bars: 100 μm. 6-8 flies per genotype. **(c-d’)** Representative ratio images of insulin-producing cells (*G9a* mutants and isogenic controls) expressing cyto-Grx1-roGFP2 specifically in these cells, and corresponding quantification in (c,c’) larvae and (d,d’) adults. *G9a* mutants show altered *E_GSH_* in insulin-producing cells during development (*p* = 0.0035) but not in adulthood (*p* = 0.16). Scale bars: 20 μm. 6-8 flies per genotype. Data are shown as individual data points with mean ±95% CI. Unpaired *t*-test. **(e)** Schematic of development-only and adult-only N-acetylcysteine amide (NACA) treatment paradigms. **(f)** Adult NACA treatment had no effect on the duration of sleep bouts in *G9a* mutants (NACA: n = 31; vehicle: n = 32) or controls (NACA: n = 30; vehicle: n = 32). **(g,g’)** Developmental NACA treatment (40 μg/mL) significantly increased sleep-bout duration and reduced sleep-bout number in *G9a* mutants (NACA-treated: n = 95; vehicle: n = 64; *p* = 0.0084), restoring sleep architecture to levels comparable to isogenic controls (vehicle: n = 63; NACA: n = 96) during the dark period (DP). (h,h’) *MsrA* knockdown in insulin-producing cells (*dilp2-Gal4>UAS*-*MsrA^RNAi-1^*, n = 39) reduced sleep-bout duration (*p* = 0.016 and *p* < 0.0001) and increased sleep-bout number (*p* = 0.0015 and *p* < 0.0001) during DP compared with both parental controls (*dilp2-Gal4/+*, n = 44; *UAS*-*MsrA^RNAi-1^/+*, n = 44). (i,i’) *MsrA* knockdown in the fat body (*UAS-Dcr2,C7-Gal4>UAS*-*MsrA^RNAi-1^*, n = 49) did not alter sleep architecture relative to parental controls (*UAS-Dcr2,C7-Gal4/+*, n = 50; *UAS*-*MsrA^RNAi-1^/+*, n = 45). Data are presented as boxplots (25^th^-75^th^ percentiles, median; whiskers indicate 5^th^-95^th^ percentiles). Kruskal-Wallis test with Dunn’s correction and Bonferroni adjustment for multiple comparisons. *p*-values are indicated as follows: \**p <* 0.05, \*\**p <* 0.01, \*\*\**p <* 0.001.

Next, we employed cytosolic Grx1-roGFP to determine whether this H_2_O_2_-dependent oxidation was accompanied by a detectable shift in the glutathione redox environment, as could be caused by the observed metabolomic alterations in methionine oxidation and cysteine biosynthesis. As glutathione is a major intracellular thiol antioxidant, its redox potential provides an integrated measure of the balance between oxidative challenge and cellular reducing capacity^44^. No significant differences were observed, at the whole-brain or fat-body level (**Fig. 5a-b’**).

Because whole-brain measurements may obscure redox changes restricted to small neuronal populations, and because G9a function in IPCs contributes to sleep consolidation, we subsequently targeted Grx1-roGFP2 specifically to IPCs. Indeed, this revealed that IPCs in *G9a* mutants exhibited increased probe oxidation during larval development (**Fig. 5c,c’**), consistent with a more oxidizing glutathione redox environment. This difference was no longer observed in adult IPCs (**Fig. 5d,d’**). We conclude that G9a mutants exhibit increased H_2_O_2_-dependent oxidation in larval brains and a developmentally restricted shift in the glutathione redox potential of IPCs.

### Developmental antioxidant treatment rescues sleep architecture in *G9a* mutants

Having established that *G9a* mutants exhibit increased H_2_O_2_ -dependent oxidation and an altered glutathione redox state, together with evidence linking redox homeostasis to sleep regulation^43,45,46^, we hypothesized that redox dysregulation in the *G9a* mutant brain contributes to the observed sleep disturbances in this KLEFS1 model. We therefore tested whether antioxidant treatment could mitigate these sleep abnormalities. We first administered the antioxidant N-acetylcysteine-amide (NACA, also known as AD4) at the previously established dose of 40 µg/ml^47,48^ to *G9a* mutants and isogenic controls. NACA is a derivative of N-acetyl-L-cysteine (NAC) in which the carboxyl group is replaced by an amide group, enhancing its membrane permeability and allowing it to efficiently cross the blood-brain barrier^49^. It acts as a precursor of both intracellular cysteine and reduced glutathione, thereby increasing ROS scavenging capacity^50^. Acute NACA treatment during adulthood did not alter sleep bout duration in *G9a* mutants or isogenic controls (**Fig. 5e,f**).

Considering that G9a function during development is critical for adult sleep architecture (**Fig. 3**), and that redox alterations were identified during this period, we next administered NACA exclusively during larval development (**Fig. 5e**). Developmental NACA treatment did not affect sleep parameters in control flies but significantly increased sleep bout duration in *G9a* mutants, fully rescuing sleep consolidation relative to control levels (**Fig. 5g,g’**). In contrast, developmental treatment with NAC, a NACA precursor with poor blood-brain permeability did not alter sleep bout duration at the therapeutic dose of 100 µM^50^ or higher (15 mM) (**Supplementary Fig. 6a-c**). Together, these results indicate that developmental redox dysregulation in the brain disrupts adult sleep integrity in the KLEFS1 model and that antioxidant treatment during this critical window can rescue the resulting sleep phenotypes.

### Reduced MsrA function in insulin-producing cells fragments nighttime sleep

What are the mechanisms underlying the developmental redox dysregulation in *G9a* mutants? Based on G9a’s histone lysine methyltransferase activity, we speculated that it may result from dysregulation of G9a target genes involved in antioxidant function. In *Drosophila*, the reduction of methionine-SO (both S and R enantiomers) back to methionine is catalyzed by the methionine sulfoxide reductase A and B (MsrA and MsrB), respectively^51^. Because we found Met-SO accumulated in *G9a* mutants (**Fig. 4**), we hypothesized that reduced methionine sulfoxide reductase activity might contribute to the observed sleep phenotype. Analysis of our previously generated mRNA-seq datasets from *G9a* null mutants^12,13^ revealed that *MsrA* expression levels tended to be lower at steady state in whole flies and were significantly decreased in heads after 6 and 12 h of exposure to paraquat, a ROS-inducing agent (**Supplementary Fig. 7a,b**).

We therefore tested whether reducing *MsrA* expression — specifically in IPCs or the FB — affects sleep architecture. IPC-specific *MsrA* knockdown using two independent RNAi constructs induced nighttime sleep fragmentation (**Fig. 5h,h’** and **Supplementary Fig. 7c,c’**), recapitulating the hallmarks of *G9a*’s sleep phenotype. In contrast, *MsrA* knockdown in the FB with the same constructs did not alter sleep architecture (**Fig. 5i,i’** and **Supplementary Fig. 7d,d’**). Unlike *MsrA*, expression of *SelR*, the *Drosophila* orthologue of *MsrB*, was not significantly reduced in *G9a* mutants (**Supplementary Fig. 8a,b**). Moreover, *SelR* knockdown in either IPCs or the FB had no effect on sleep (**Supplementary Fig. 8c-f’**). Together, our findings demonstrate that *MsrA* activity in IPCs contributes to nighttime sleep consolidation and argue that reduced expression of this enzyme contributes to the sleep disturbances observed in *G9a* mutants, preventing regeneration of reduced methionine in these cells.

### Sleep disturbances in KLFS1 models can be behaviorally rescued with sleep-restriction therapy

Last, because the necessity of antioxidant treatment during development will be limiting clinical applicability of our findings, we also asked whether a behavioral intervention could improve sleep quality in our preclinical KLEFS1 models. Sleep restriction therapy (SRT) is a key component of cognitive behavioral therapy for insomnia (CBT-I), the first-line intervention in humans to treat chronic insomnia^52^. This non-invasive and non-hazardous approach enhances sleep drive by restricting the opportunity for sleep, leading to more efficient and consolidated sleep. In *Drosophila,* sleep opportunity can be restricted by shortening the duration of the darkness period to which the flies are entrained^53^. We have recently reported that an adult SRT regime reversed sleep fragmentation of developmental origin in kismet, a fly model of two related neurodevelopmental disorders^19^. Therefore, we tested whether SRT may also be able to counteract sleep fragmentation in the KLEFS1 model.

First, we restricted the opportunity for sleep in *G9a* mutants by reducing the duration of the dark period to 10 hours (14:10 LD cycle) and evaluated sleep efficiency, a value utilized in the clinic to score sleep quality. In *Drosophila,* in analogy to humans, it represents the time spent asleep in the dark sleep opportunity window. The 14:10 LD SRT regime only insignificantly increased sleep efficiency as well as bout duration, yet significantly decreased bout number in *G9a* mutants (**Supplementary Fig. S9a-e**). Restricting the dark period further to only 8 hours (16:8 LD cycle, **Fig. 6a**) decreased total sleep time during this shorter, relative night of mutants kept under SRT conditions (**Supplementary Fig. 9f**), however it fully rescued the lower sleep efficiency of *G9a* mutants (**Fig. 6b**). Moreover, sleep bout duration and sleep bout number were normalized, with neither parameter differing significantly from controls entrained at 12:12 LD (**Fig. 6c,d**). Wake after sleep onset, another parameter analogous to the clinical assessment, increased in the majority of investigated KLEFS1 individuals (**Table 1**), was also rescued (**Fig. 6e**). Taken together, our data show that SRT can improve the disturbed sleep architecture caused by loss of G9a.

**Figure 6.**
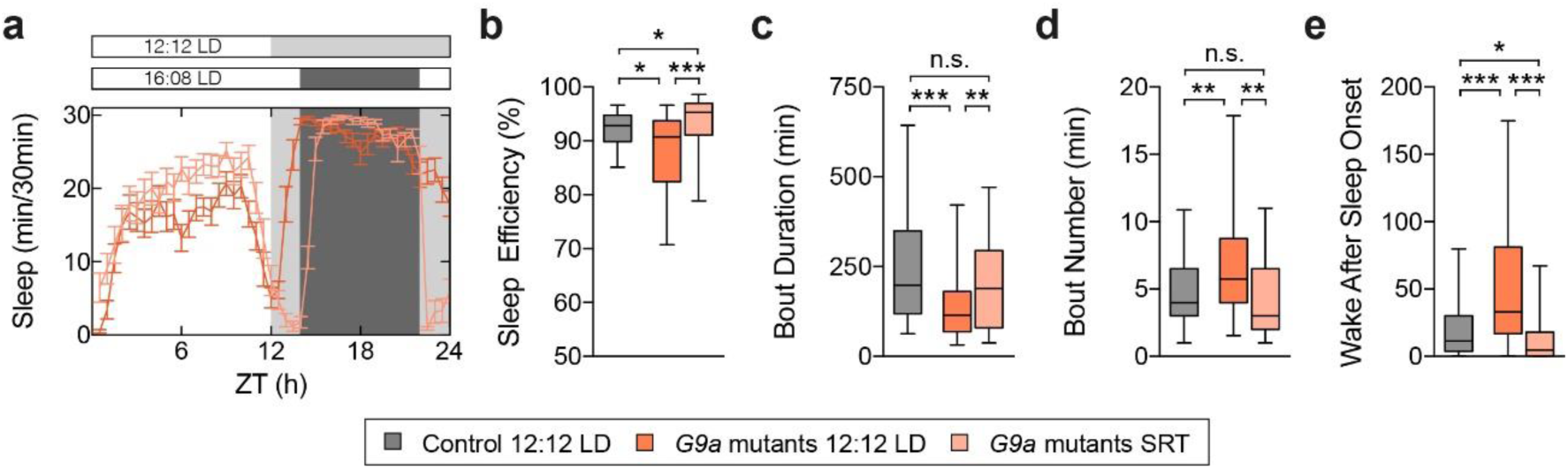
Sleep-restriction therapy rescues sleep architecture in *Drosophila* G9a models. **(a)** Representative sleep profiles of *G9a* mutant flies entrained either at 12:12 LD (= 21) or 16:8 LD (shortened dark period ZT14-22, n = 22). **(b)** Quantification of sleep efficiency, **(c)** sleep bout duration and **(d)** bout number, and **(e)** WASO of *G9a* mutant flies entrained either at 12:12 LD (n = 71) or 16:8 LD (n = 69) compared to isogenic controls (n = 72) during the DP. SRT successfully improved sleep efficiency and architecture in *G9a* mutant flies. Data are represented as boxplots that extend from the 25^th^ to the 75^th^ percentiles with the median indicated. Whiskers indicate the 5^th^ and 95^th^ percentiles. Kruskal-Wallis test with Dunn’s correction. Bonferroni correction for multiple testing. *p*-values are indicated as follows: \**p <* 0.05, \*\**p <* 0.01, \*\*\**p <* 0.001.

## Discussion

Our study moved beyond the recognition that sleep disturbances are common in the neurodevelopmental disorder KLEFS1^7,9,10,54–56^, providing a dedicated characterization of the clinical sleep phenotype through caregiver questionnaires and sleep diaries. Although caregiver-reported sleep diaries are likely to quantitatively underestimate the frequency of nighttime awakenings, the patterns observed across participants and the two approaches we utilized here generated highly consistent data demonstrating that deficits in sleep maintenance leading to night awakening and sleep fragmentation are the predominant hallmark of sleep disturbances in KLEFS1. More quantitative approaches, including actigraphy or video-based monitoring, will be important to refine the frequency and duration of these awakenings more precisely, though this may methodologically be challenging among this population. Further monitoring these awakenings and recognizing them early on is particularly relevant given that severe sleep disturbances are frequently reported around regressive periods in KLEFS1 and are already recognized clinically as an early feature of these episodes^4,9^. Whether sleep disruption contributes to the subsequent deterioration in adaptive functioning, rather than simply accompanying regressions, remains an important open question. Finally, the association between common variation at the *EHMT1* locus and sleep-related traits in the general populations extends these observations beyond KLEFS1, indicating that EHMT1 is not only a prerequisite for brain development supporting healthy sleep, but is also involved in its physiological regulation. Common variation at the intact *EHMT1* copy in individuals with KLEFS1 is, based on our findings, thus also likely to contribute to the clinical variability of sleep disturbances in individuals with KLEFS1.

To determine how EHMT1/G9a disruption causes sleep disturbances, we took advantage of *Drosophila —* an already established model to study mechanisms in sleep and neurodevelopmental disorders — and its sophisticated genetic toolbox. Our work demonstrated the EHMT1/G9a family of chromatin remodellers as conserved regulators of sleep maintenance. Thus, sleep joined a growing list of metabolic, cognitive, and behavioral phenotypes shared between *Drosophila* models of EHMT1/G9a dysfunction and individuals with KLEFS1, including altered lipid metabolism, inflammation, learning and memory, and habituation^4,11,13,14^. This convergence further supports the value of *Drosophila* as a clinically relevant model for investigating the molecular mechanisms underlying KLEFS1 phenotypes and for identifying potential therapeutic strategies.

Cell-specific and temporal analyses identified the IPCs and the developing FB as critical tissue in the KLEFS1 sleep pathology, pointing to systemic metabolic regulation as an important mediator of the sleep phenotype. This is of particular interest in KLEFS1, where metabolic abnormalities, including overweight and obesity, are also common^4,7^, and consistent with the established links between insulin signaling, energy homeostasis, and sleep^57^. Our metabolomics and redox sensor analyses as well as genetic (MsrA) and pharmacological (antioxidant) intervention experiments provide multiple lines of evidence for increased ROS in the developing *G9a* brain and in IPCs lying at the core of the observed sleep fragmentation. This KLEFS1-specific finding is moreover in line with an established bidirectional relationship between ROS accumulation and sleep^32,43^. Together, with our previous work demonstrating a role for G9a in energy metabolism and stress tolerance^13,14^, and our recent work linking metabolic dysregulation to developmental regression in KLEFS1^58^, these findings suggest that impaired metabolic homeostasis may represent a broader vulnerability of EHMT1/G9a dysfunction, particularly under physiological stress.

Finally, sleep (opportunity) restriction therapy improved sleep consolidation in *G9a* mutants, providing further proof-of-principle that sleep disturbances arising from developmental metabolic dysfunction can be ameliorated through acute behavioral intervention. This is a highly potent direction for future clinical investigation. The methodology is standardly applied in sleep clinics for the treatment of insomnia but has so far been hardly considered for individuals with NDDs. Whereas improving sleep of individuals with NDDs promises to readily lift up the quality of life of caretakers and families, it may - given the broad importance of sleep – have benefits for cognitive and many other physiological functions^59–61^.

Altogether, our cross-species study established EHMT1/G9a as an evolutionarily conserved regulator of sleep integrity and supports a model in which developmental disruption of metabolic and redox homeostasis leads to persistent sleep fragmentation. In addition to providing new perspectives on the developmental origins of sleep dysfunction, our findings highlight the modifiability of the resulting adult sleep phenotype, providing a foundation for mechanistically- and clinically-grounded investigations of sleep disturbances in KLEFS1.

## Materials and methods

### Sleep assessment in individuals with KLEFS1

After informed consent, parents of patients agreed to participate and were asked to complete the Modified Simonds & Parraga Sleep Questionnaire^19–21^ digitally using the survey package from Castor EDC. Additionally, they were asked to fill a graphical sleep diary for two consecutive weeks^19–21^. Dutch participants filled it in Dutch and on paper and international cohort in English directly online.

Sleep parameters were calculated from the sleep diaries using the average of each individual sleep indices. The following parameters were assessed with a 15-minute bin resolution based on parent-reported sleep-wake calendars: time in bed, total sleep time (TST), sleep onset latency (SOL), wake after sleep onset (WASO), number of night awakenings, and sleep efficiency. TIB and SOL were calculated from the start of sleep opportunity (*i.e.,* when the light was turned off). Sleep efficiency was calculated as TST/time in bed × 100 and was considered deviant if below 85%. SOL was defined as the time between going to bed and sleep onset. WASO was calculated as the total time awake between sleep onset and time out of bed. Nighttime awakenings due to being awakened by a caretaker or bed wetting incidents were not included into the WASO calculation nor in the number of awakenings. Reference values for typically developing children and adolescents were taken from previous studies^62,63^.

However, based on the International Classification of Sleep Disorders 3 (ICSD-3) criteria, which define a sleep disorder as complaints occurring at least three times per week, we used a threshold of three or more nights per week. Particular days were excluded from analysis when the participants’ sleep was affected by sickness or if caregivers forgot to fill the diary.

### Gene-based associations of *EHMT1*

On June 28, 2021, we retrieved gene-based *P*-values for *EHMT1* (NCBI Gene ID: 79813) for all available GWAS (∼4,000) in the GWAS atlas^23^ for sleep related traits (GWAS n = 11). We queried GWAS results from the sub-chapters “Sleep Functions” (Domain: Psychiatric; Chapter: Mental Functions) and “Sleep Disorders” (Domain: Neurological; Chapter: Diseases of the Nervous System). If duplicated traits were identified, we selected the GWAS including the largest sample size and the GWAS that used combined data from males and females. In GWAS atlas, for the gene-based analyses, MAGMA v1.06^64^ was used. MAGMA gene analysis was performed using 19,436 protein coding genes obtained from biomaRt (primary ID is Ensembl ID v92 GRCh37) which are mapped to entrez ID from NCBI. SNPs are assigned to genes with a 1 Kb window on each side. For all gene-based analyses, the default model, SNP-wide (mean) was used.

### *Drosophila* stocks and husbandry

Flies were reared on standard medium containing yeast, cornmeal, agar, and sugar at 25°C and 60% humidity. For experiments, male flies were entrained in a 12:12 LD cycle, unless indicated otherwise The following *Drosophila* stocks were used for this study: *w^+^* wildtype, *G9a* mutants [*G9a^DD1^* described in ^11^], the genetic background control for Vienna *Drosophila* Resource Center (VDRC) GD library (VDRC #60000), *UAS-G9a^RNAi^*(VDRC #25474), genetic background control for VDRC KK library (VDRC #60100), genetic background control for attP2 TRiP library [Bloomington *Drosophila* Stock Center (BDSC) #36303], *R14H06*-*Gal4* (BDSC #48667), *R96A08-Gal4* (BDSC #48030), *dilp2-Gal4/CyO (BDSC #37516)*, *C7-Gal4;UAS-Dcr2 (*kindly provided by Marek Jindra^65^), *FB-Gal4* (kindly provided by Tomáš Doležal), *UAS-cyto-Grx1-roGFP2* (BDSC #67662), *UAS-cyto-Grx1-roGFP2;tub-Gal4* (BDSC #67663), *tub-cyto-roGFP2-Orp1* (BDSC #67670), *UAS-MsrA^RNAi-1^* (BDSC #42877), *UAS-MsrA^RNAi-2^* (VDRC # 26009), *UAS-SelR^RNAi-1^* (VDRC #110755), *UAS-SelR^RNAi-2^* (VDRC #25999).

### Sleep monitoring and analysis

The locomotor activity and sleep patterns were tracked using the *Drosophila* Activity Monitor (DAM2) system developed by Trikinetics (Waltham, MA, USA). In brief, 4-to-7-day old male flies aged (unless otherwise stated) were individually transferred to glass tubes (65mm x 5 mm) containing standard food, without the use of CO_2_ anesthesia. These tubes were then loaded into the DAM2 systems, and then flies were allowed to acclimate to both the activity monitors and the food conditions for a minimum of 12 hours. Subsequently, they were monitored for four days at a temperature of 25°C under a 12:12 light:dark cycle, unless otherwise stated.

In brief, 4- to 7-day-old male flies (unless specified otherwise) were individually transferred without CO_2_ anesthesia to glass tubes (65mm x 5 mm) containing standard food, loaded into the DAM systems, allowed to acclimatize to activity monitors and food for at least 12h and monitored for four days at 25°C in a 12:12 LD cycle unless specified otherwise. The motion of the flies was detected using infrared light beams integrated into the monitors. To quantify activity and sleep (defined in *Drosophila* as periods of five or more minutes of inactivity)^66,67^, specific parameters were extracted using the publicly available Sleep and Circadian Analysis MATLAB Program (SCAMP)^68^ designed for MATLAB. The sleep data presented in this study represents the average over the four-day data acquisition period, unless specified otherwise.

### Sleep deprivation

Mechanical sleep deprivation was accomplished as previously described^16^. In essence, flies in DAM2 monitors were placed on a Trikinetics vortexer mounting plate (Waltham, MA, USA) and their activity was recorded for three consecutive days. On the second day, they were shaken for 2 seconds randomly within every 20-second window for 12 hours during the dark period (ZT 12-24). The capability of each tested genotype to rebound was assessed by comparing the total sleep during ZT 0-3 in the preceding day in unperturbed conditions (baseline) and the same period following sleep deprivation (recovery).

### Circadian analysis

Circadian analysis was performed as previously described^16^. In brief, flies were loaded into the DAM system 4-7 days after eclosion as described above and entrained to a 12:12 LD cycle for 3 days before being transferred to constant darkness (DD). Locomotor activity during days 2-7 in DD was analyzed in the software Clocklab (Actimetrics, Wilmette, IL). Fast Fourier Transform (FFT)^69^ was performed and the maximum amplitude of the FFT was calculated and compared across genotypes. According to their FFT, flies were categorized as strongly rhythmic (FFT ≥ 0.05), moderately rhythmic (0.05 > FFT ≥ 0.03), weakly rhythmic (0.03 > FFT ≥ 0.01), or arrhythmic (FFT < 0.01).

### Sleep Restriction Therapy

4- to 7-day-old male flies were loaded into the DAM systems as described above, allowed to acclimatize to activity monitors and food for at least 12h and monitored for four days at 25°C in either a 12:12 LD, 14:10 LD or 16:08 LD cycle. Sleep data shown is the average of the last two days of monitoring.

### Sample collection and processing for metabolomics analysis

4- to 7-day-old male *G9a* mutant flies and respective isogenic controls entrained in a 12:12 LD cycle were frozen by immersion in liquid nitrogen. Four replicates per genotype, each consisting of 100 heads, were collected in parallel, stored at –80°C and sent to Metabolon Inc. (Morrisville, NC) for sample processing.

### Metabolomics sample preparation and analysis (Metabolon Inc.)

Samples for Ultrahigh Performance Liquid Chromatography-Tandem Mass Spectroscopy (UPLC-MS/MS) were prepared using the automated MicroLab STAR® system from Hamilton Company. Several recovery standards were added prior to the first step in the extraction process for quality control purposes. To remove protein, dissociate small molecules bound to protein or trapped in the precipitated protein matrix, and to recover chemically diverse metabolites, proteins were precipitated with methanol under vigorous shaking for 2 min (Glen Mills GenoGrinder 2000) followed by centrifugation. The resulting extract was divided into five fractions: two for analysis by two separate reverse phase (RP)/UPLC-MS/MS methods with positive ion mode electrospray ionization (ESI), one for analysis by RP/UPLC-MS/MS with negative ion mode ESI, one for analysis by HILIC/UPLC-MS/MS with negative ion mode ESI, and one sample was reserved for backup. Samples were placed briefly on a TurboVap® (Zymark) to remove the organic solvent. The sample extracts were stored overnight under nitrogen before preparation for analysis. All methods utilized a Waters ACQUITY ultra-performance liquid chromatography (UPLC) and a Thermo Scientific Q-Exactive high resolution/accurate mass spectrometer interfaced with a heated electrospray ionization (HESI-II) source and Orbitrap mass analyzer operated at 35,000 mass resolution. The sample extract was dried then reconstituted in solvents compatible to each of the four methods. Each reconstitution solvent contained a series of standards at fixed concentrations to ensure injection and chromatographic consistency. One aliquot was analyzed using acidic positive ion conditions, chromatographically optimized for more hydrophilic compounds. In this method, the extract was gradient eluted from a C18 column (Waters UPLC BEH C18-2.1×100 mm, 1.7 µm) using water and methanol, containing 0.05% perfluoropentanoic acid (PFPA) and 0.1% formic acid (FA). Another aliquot was also analyzed using acidic positive ion conditions, however it was chromatographically optimized for more hydrophobic compounds. In this method, the extract was gradient eluted from the same afore mentioned C18 column using methanol, acetonitrile, water, 0.05% PFPA and 0.01% FA and was operated at an overall higher organic content. Another aliquot was analyzed using basic negative ion optimized conditions using a separate dedicated C18 column. The basic extracts were gradient eluted from the column using methanol and water, however with 6.5 mM Ammonium Bicarbonate at pH 8. The fourth aliquot was analyzed via negative ionization following elution from a HILIC column (Waters UPLC BEH Amide 2.1×150 mm, 1.7 µm) using a gradient consisting of water and acetonitrile with 10mM Ammonium Formate, pH 10.8. The MS analysis alternated between MS and data-dependent MS scans using dynamic exclusion. The scan range varied slighted between methods but covered 70-1000 m/z. Raw data was extracted, peak-identified and QC processed using Metabolon’s hardware and software. Compounds were identified by comparison to Metabolon’s library entries of purified standards or recurrent unknown entities.

A variety of curation procedures were carried out to remove system artifacts, mis-assignments, and background noise. Metabolon data analysts use proprietary visualization and interpretation software to confirm the consistency of peak identification among the various samples. Library matches for each compound were checked for each sample and corrected if necessary. Peaks were quantified using area-under-the-curve.

Metabolite abundance data were normalized using probabilistic quotient normalization (PQN), followed by generalized logarithm (glog) transformation and unit variance (UV) scaling. Then, a robust linear regression model was fitted to the data using the “lmFit” function from the "limma" R package^69^, with method = "robust" to minimize the influence of potential outlying samples. Finally, an empirical Bayes method was applied using the “eBayes” function by borrowing information across metabolites. P-values were adjusted for multiple testing using the false discovery rate (FDR) method, with an FDR <0.1 considered statistically significant. The functional relevance of differentially abundant metabolites was further investigated through over-representation analysis based on the Kyoto Encyclopedia of Genes and Genomes (KEGG) database. Pathway enrichment significance was assessed using a hypergeometric test, and P-values were corrected for multiple testing using the Storey method. Statistical significance was set at <0.1.

### Probing ROS with genetic redox sensors

Brains from third-instar (L3) larvae and 3-day-old adult flies, as well as fat bodies from L3 larvae, were dissected in chilled PBS containing 20 mM N-ethyl maleimide (NEM). Tissues were fixed within 20 minutes of dissection in 2% paraformaldehyde (PFA) and 20 mM NEM for 1 hour at room temperature^70^. Following fixation, samples were washed in PBS, mounted immediately on glass slides in Vectashield H-1000, and sealed. Slides were imaged within two days. Confocal images were acquired using a Zeiss LSM 880 microscope. Sequential excitation at 405 nm and 488 nm was used, and emission was collected between 495–525 nm.

Image analysis followed established methods ^44,70^. Background pixels were first removed by applying a mask generated using an auto-threshold and a 3-pixel radius median filter. A ratio image was then produced by dividing the filtered 405 nm channel by the filtered 488 nm channel on a pixel-wise basis. To further eliminate debris and autofluorescent tracheal signal, pixels with a 405:488 nm ratio greater than 4:1 were excluded. For whole-brain mounts, trachea were manually removed from each optical section prior to generating a maximum-intensity projection. Mean gray values of the final images were recorded, and images were visualized using the “Fire” lookup table.

### Antioxidant feeding

*N-*acetylcysteine amide (NACA; Torcis Bioscience, #5619) was incorporated into fly food at 50°C to a final concentration of 250 µM. The concentration we selected based on previous studies^47,48^. For developmental treatment, NACA was added to the food on which flies were maintained from egg laying until eclosion. Newly eclosed adults were subsequently transferred to untreated food before and during experiments. For adult-only treatment, newly eclosed flies were instead transferred and maintained on food containing NACA before and during experiments.

For comparison, *N-*acetyl-L-cysteine (NAC; Sigma-Aldrich, #A7250) was incorporated into food at 50°C to a final concentration of 100 µM^50^ or 15 mM. Flies were reared on food containing NAC during development, and after eclosion, were transferred to untreated food before and during experiments. Antioxidant food was prepared fresh the day before experiments and stored at 4°C until experiments.

### Statistical Analysis

Adult *Drosophila* sleep data statistical analysis was carried out in GraphPad Prism version 7 (GraphPad Software, San Diego, California, USA). Datasets comprised by two groups that followed a Gaussian distribution but showed differences in their standard deviation with the controls were analyzed with a two-tailed unpaired Welch’s t-test. For groups of more than two genotypes, Kruskal-Wallis test with Dunn’s multiple comparisons test was performed to assess significance between the tested groups. *Post hoc* Bonferroni correction for multiple testing was further applied for the number of tests performed on a dataset per genotype to determine the corrected two-sided significance level. Only *p*-values that pass the corrected significance level are indicated in the figures. All data shown is from at least three independent experiments (n = 3) unless specified otherwise. For sleep deprivation experiments, statistical significance was determined with a two-tailed paired t-test between baseline and recovery day per genotype*. Post hoc* Bonferroni correction for multiple testing was applied to correct for the number of tests performed on a dataset per genotype. Only *p*-values that withstand multiple testing correction are indicated in the figures. Three independent biological replicates were performed. For the metabolomics data analysis, standard statistical analyses were performed in ArrayStudio. Significance was estimated with a two-tailed unpaired Welch’s t-test.

## Data Availability

Data reported in the current study may be shared by the lead contact upon reasonable request.

## Supplementary Figures

**Supplementary Figure 1.**
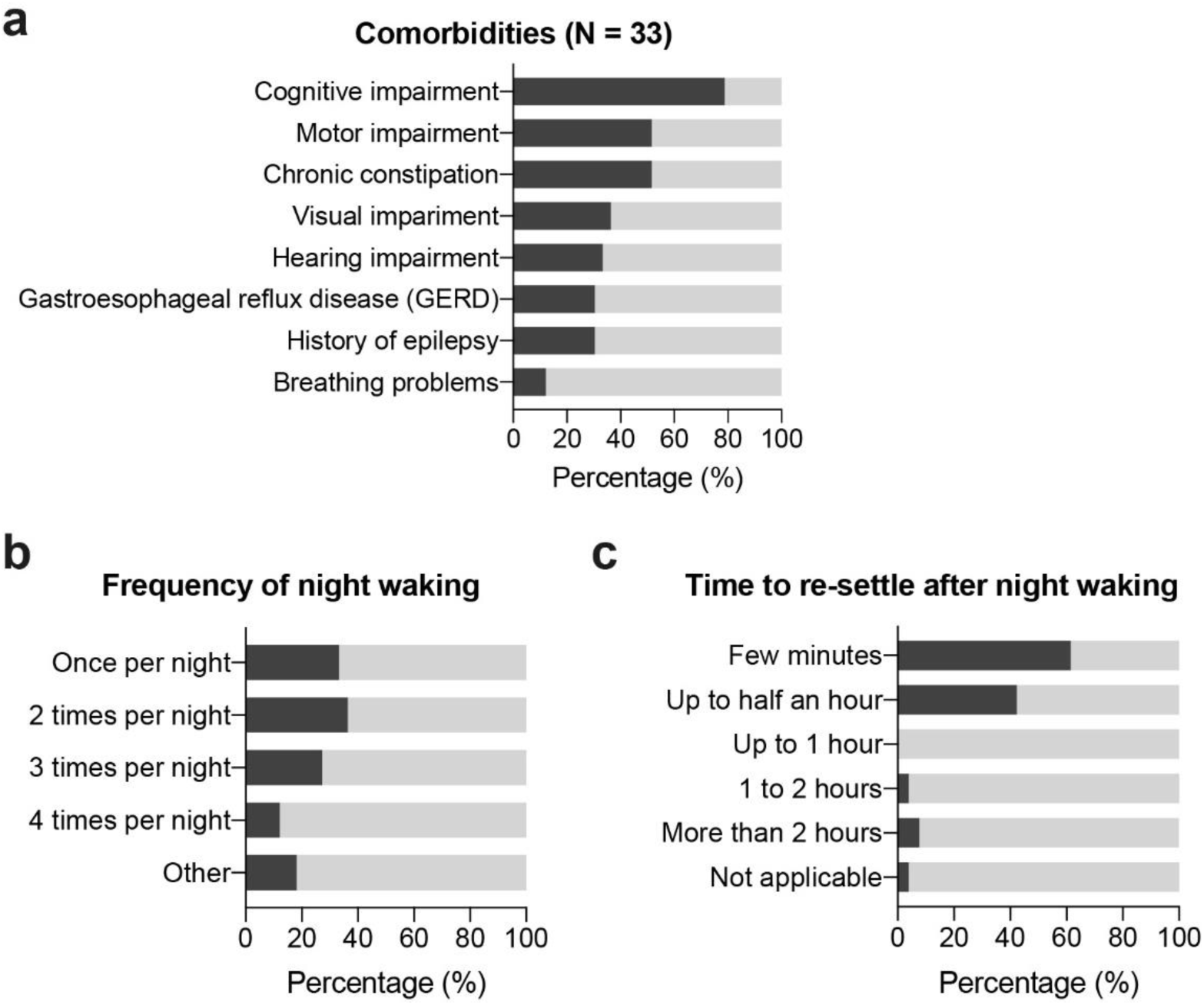
KLEFS1 cohort comorbidities, frequency of night waking, and time to resettle after waking. **(a)** Comorbidities present in our KLEFS1 cohort. **(b)** Distribution of the frequency of night wakings and **(c)** the time it takes caregivers to resettle their child after they wake up during the night in our KLEFS1 cohort.

**Supplementary Figure 2.**
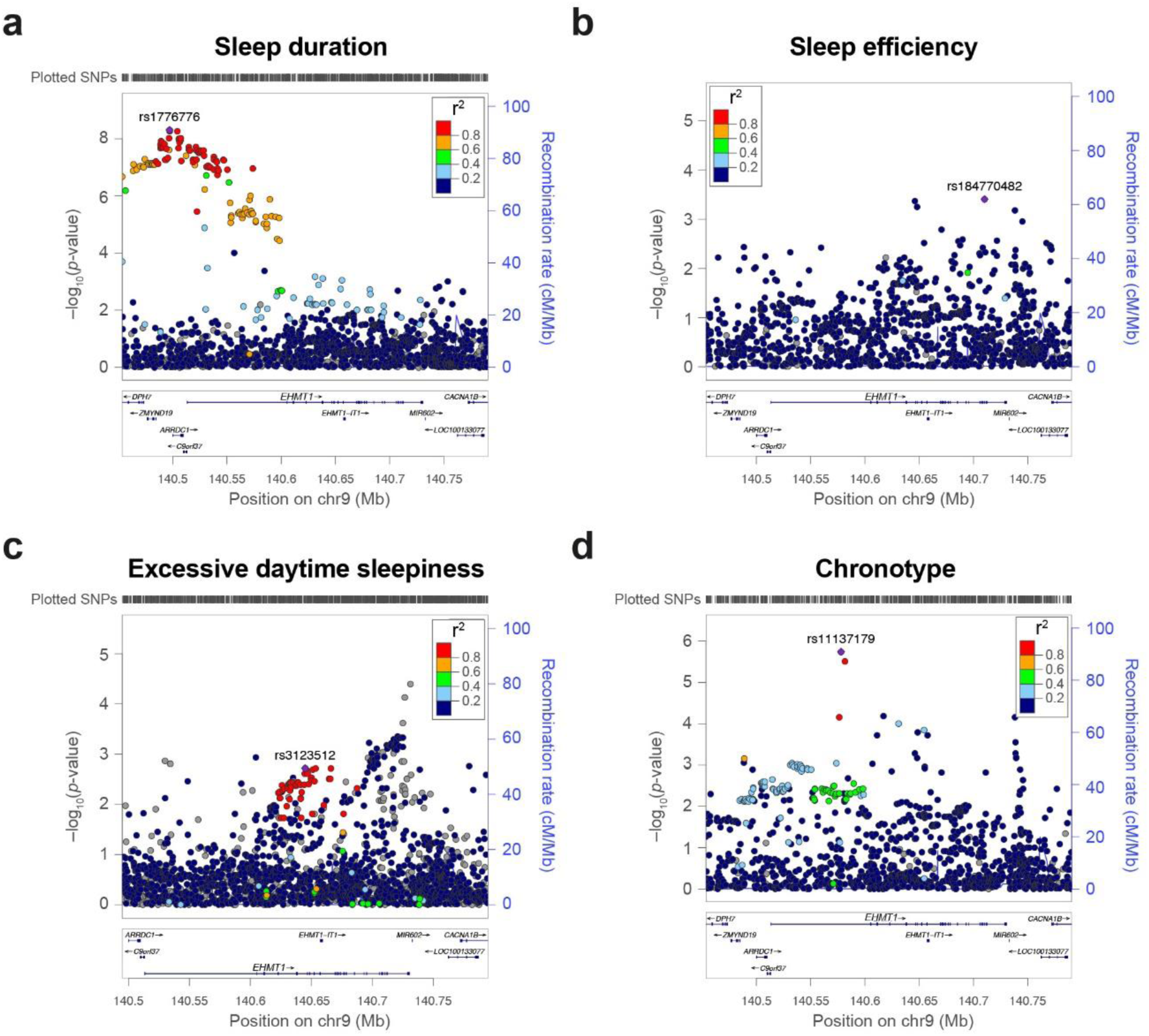
Common variation in *EHMT1* is associated with sleep duration, sleep efficiency, excessive daytime sleepiness, and chronotype. **(a)** Regional association plots showing association signals for sleep duration, **(b)** sleep efficiency, **(c)** excessive daytime sleepiness, and **(d)** chronotype at the *EHMT1* locus. Data are shown as −log_10_(*p*-value) for individual SNPs. The color of each marker reflects its linkage disequilibrium (r^2^) with the strongest associated SNP indicated as a purple diamond. The recombination rate is indicated in blue. Chr, chromosome; cM, centimorgan; Mb, megabase.

**Supplementary Figure 3.**
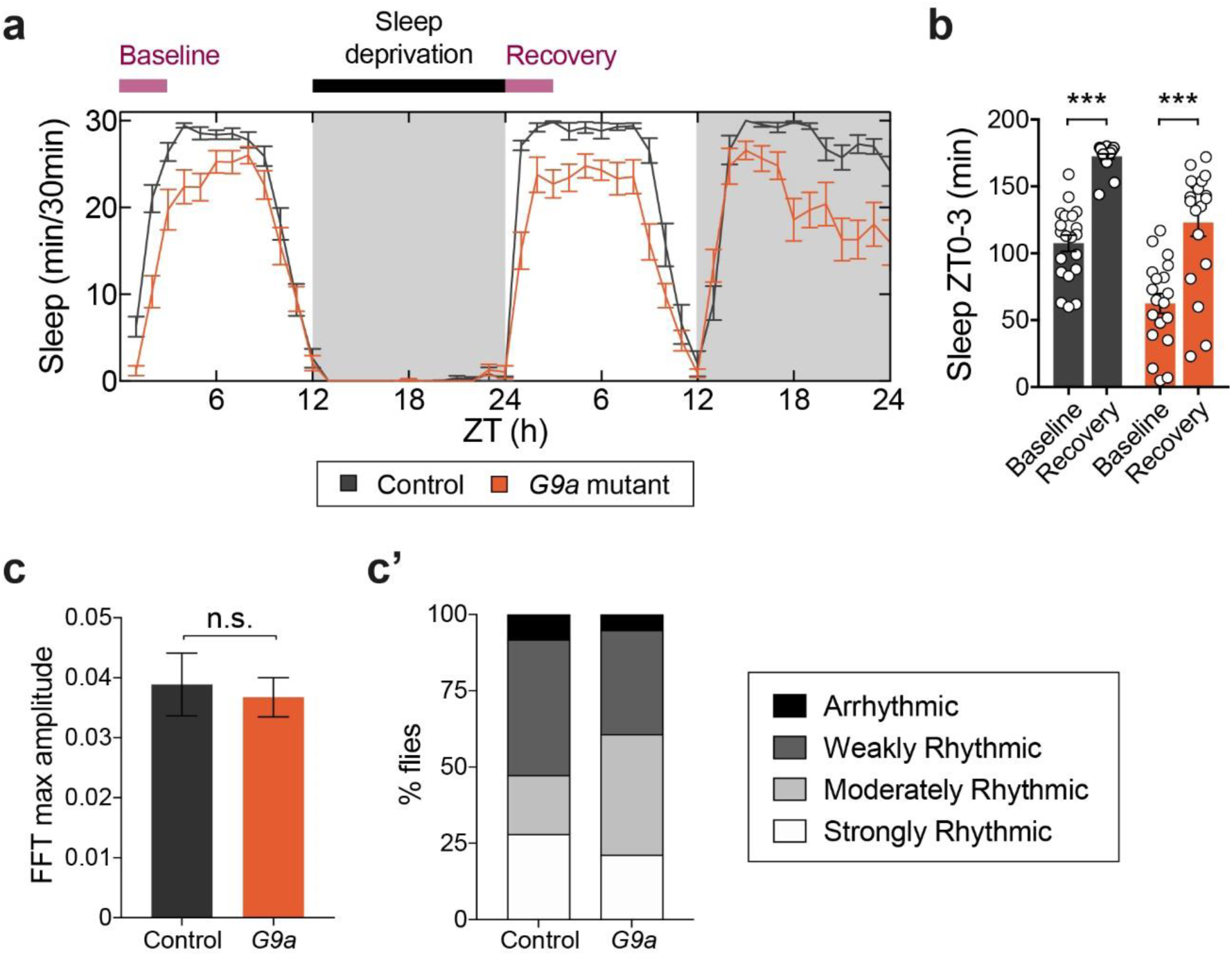
*G9a* mutants have intact homeostatic rebound after sleep deprivation and maintain rhythmicity in free-running. **(a)** Representative sleep profile of flies at ZT0-3 before (baseline) and after (recovery) 12h mechanical sleep deprivation. **(b)** Both isogenic controls (in black, n = 20) and *G9a* mutant flies (in orange, n = 19) showed significant rebound after 12h of mechanical sleep deprivation (both groups *p <* 0.0001). Two-tailed paired Students *t*-test. Bonferroni correction for multiple testing. **(c)** Comparison of rhythm strength in *G9a* mutants and their isogenic controls as measured by Fourier transform (FFT). Two-tailed unpaired Welch’s *t* test (*p* = 0.73). **(c’)** Proportion of strongly rhythmic (FFT ≥ 0.05), moderately rhythmic (0.05 > FFT ≥ 0.03), weakly rhythmic (0.03 > FFT ≥ 0.01), and arrhythmic flies (FFT < 0.01). Data are represented as mean ± SEM. *p*-values are indicated as follows: \*\*\**p <* 0.001.

**Supplementary Figure 4.**
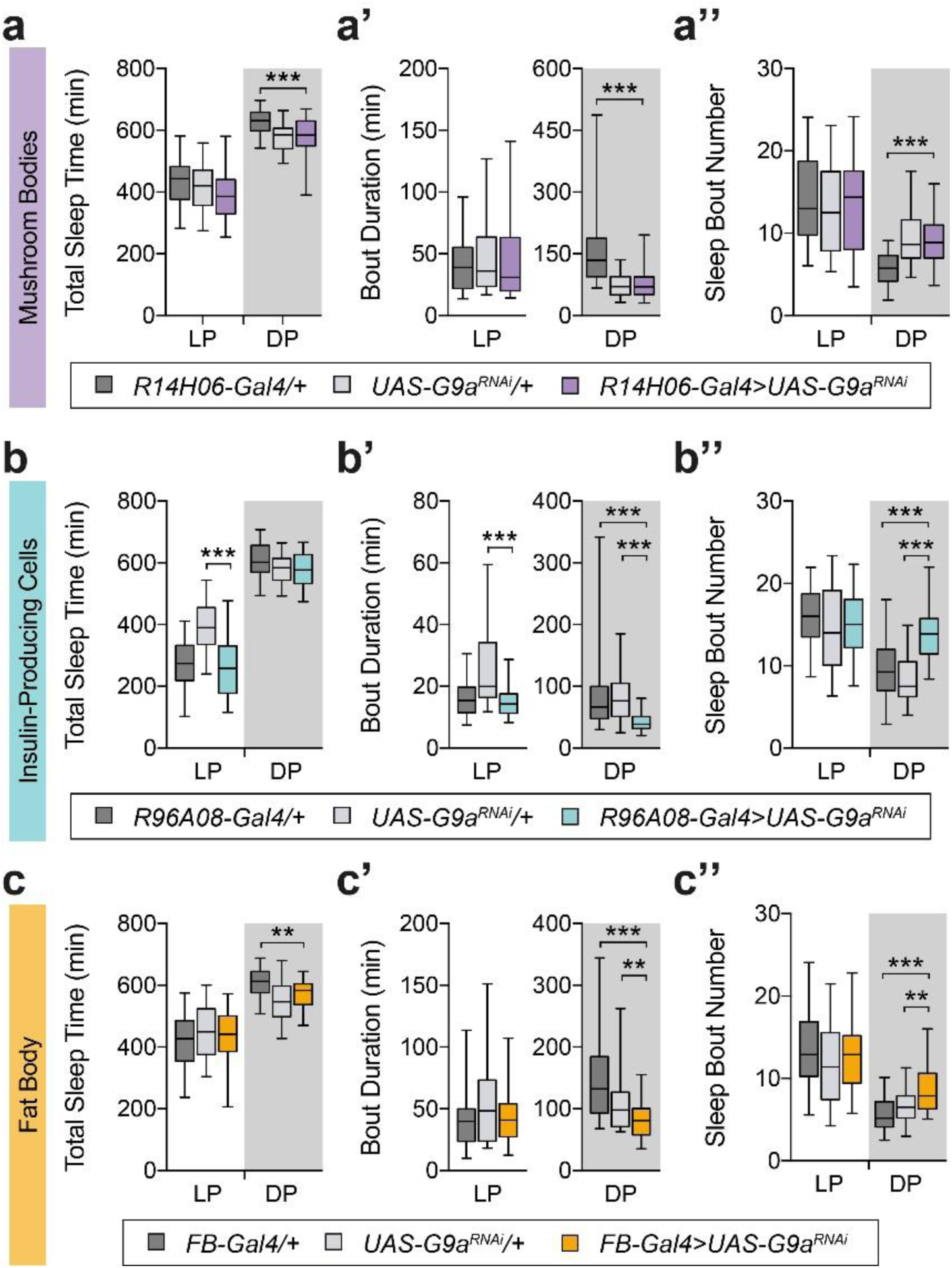
G9a does not regulate sleep in mushroom body neurons. *G9a* knockdown with independent insulin-producing cells and fat body drivers recapitulate sleep fragmentation. **(a-c)** Total sleep time duration, **(a’-c’)** average sleep bout duration and **(a’’-c’’)** number of those sleep bouts in the light period (LP, ZT0-12) and dark period (DP, ZT12-24). **(a-a’’)** *G9a* knockdown in mushroom body neurons (*R14H06*-*Gal4 > UAS-G9a^RNAi^*, n = 70) does not alter sleep duration nor architecture when compared to both compared to both parental controls (*R14H06*-*Gal4/+*, n = 69; *UAS*-*G9a^RNAi^/+*, n = 72). **(b-b’’)** *G9a* knockdown flies in insulin-producing cells neurons (*R96A08-Gal4 > UAS*-*G9a^RNAi^*, n = 70) leads to shorter sleep bouts (*p* < 0.0001) with an increase in their number (*p* < 0.0001) exclusively during DP compared to both parental controls (*R96A08-Gal4/+*, n = 71; *UAS*-*G9a^RNAi^/+*, n = 72). **(c-c’’)** *G9a* knockdown flies in the fat body (*FB-Gal4 > UAS*-*G9a^RNAi^*, n = 44) leads to shorter sleep bouts (*p* < 0.0001 and *p* = 0.008) with an increase in their number (*p* < 0.0001 and *p* = 0.006) exclusively during the DP compared to both parental controls (*FB-Gal4/+*, n = 44; *UAS*-*G9a^RNAi^/+*, n = 38). Data are represented as boxplots that extend from the 25^th^ to the 75^th^ percentiles, with the median indicated. Whiskers indicate the 5^th^ and 95^th^ percentiles. One-way ANOVA with Bonferroni correction for normally distributed data or Kruskal-Wallis test with Dunn’s correction. Bonferroni correction for multiple testing. *p*–values are indicated as follows: \**p <* 0.05, \*\**p <* 0.01, \*\*\**p <* 0.001.

**Supplementary Figure 5.**
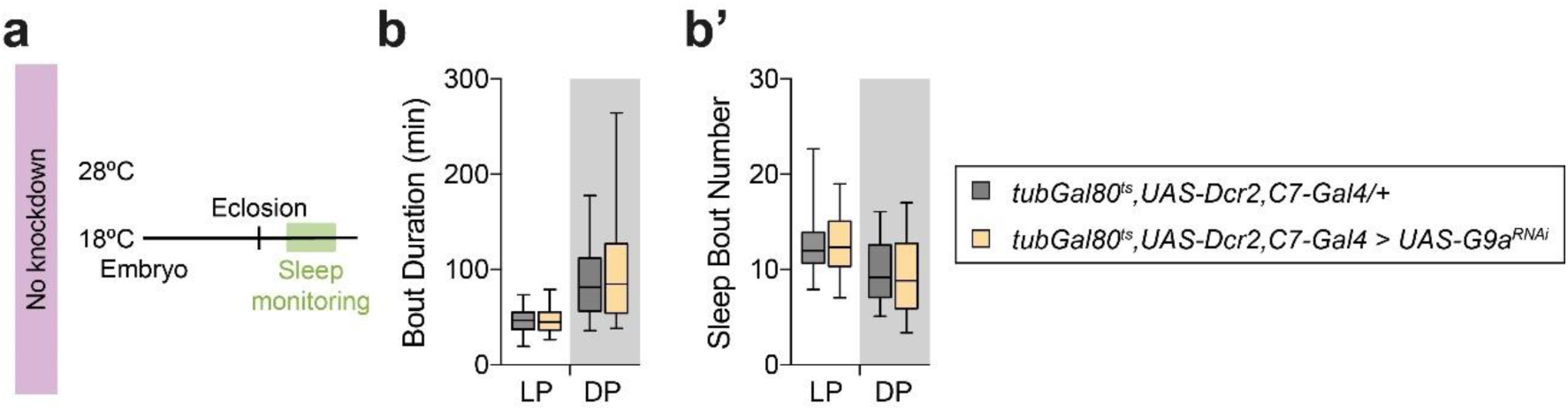
TARGET *G9a* knockdown flies do not show fragmented sleep when reared at a restrictive temperature. **(a)** Temperature scheme used to induce control for temporal pre- and post-eclosion *G9a* knockdown, resulting in no *G9a* knockdown. **(b)** Average sleep bout duration and **(b’)** number of sleep bouts in flies when *G9a* knockdown in the fat body is not induced as flies are kept at 18°C (*tub-Gal80^ts^, C7-Gal4, UAS-Dcr2 > UAS*-*G9a^RNAi^*, n = 62) compared to isogenic controls (*tub-Gal80^ts^, C7-Gal4, UAS-Dcr2/+*, n = 44). Data are represented as boxplots that extend from the 25^th^ to the 75^th^ percentiles with the median indicated. Whiskers indicate the 5^th^ and 95^th^ percentiles. Two-tailed unpaired t-test or Mann-Whitney U test. Bonferroni correction for multiple testing.

**Supplementary Figure 6.**
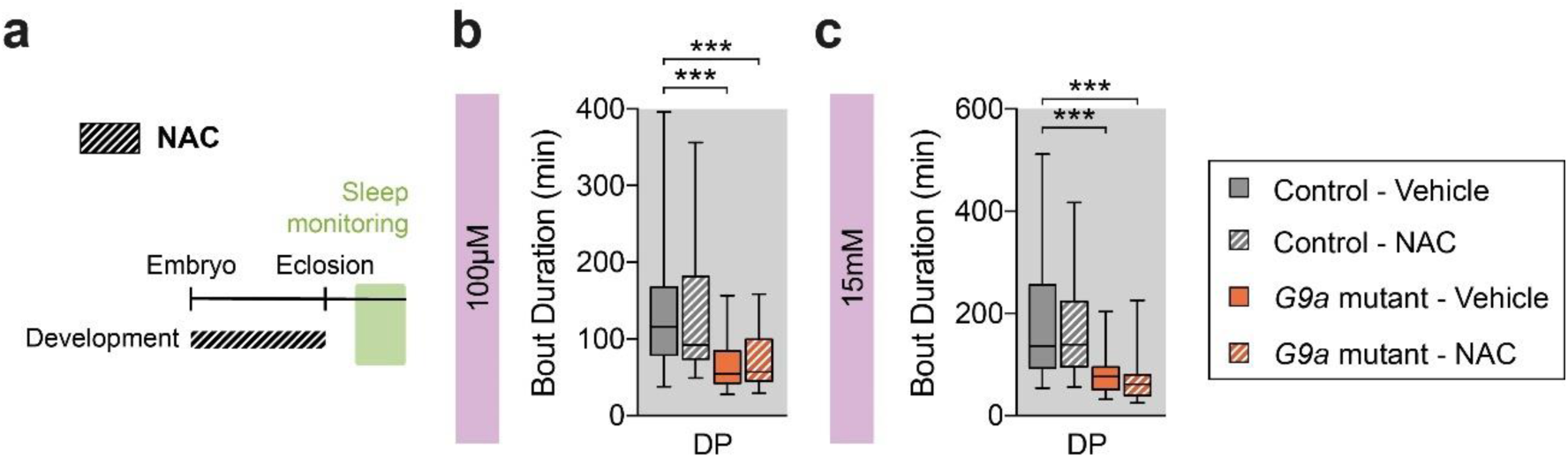
NAC treatment does not rescue nighttime sleep fragmentation. **(a)** Schematic of development-only *N*-acetyl-L-cysteine (NAC) treatment paradigm. **(b,c)** Developmental NAC treatment had no effect on the duration of sleep bouts at **(b)** 100 µM in *G9a* mutants (NAC: n = 63; vehicle: n = 64) or controls (NAC: n = 63; vehicle: n = 64), or **(c)** 15 mM in *G9a* mutants (NAC: n = 54; vehicle: n = 62) or controls (NAC: n = 63; vehicle: n = 60). Data are represented as boxplots (25^th^-75^th^ percentiles, median; whiskers indicate 5^th^-95^th^ percentiles). Kruskal-Wallis test with Dunn’s correction and Bonferroni adjustment for multiple comparisons. DP: dark period. *p*-values are indicated as follows: \**p <* 0.05, \*\**p <* 0.01, \*\*\**p <* 0.001.

**Supplementary Figure 7.**
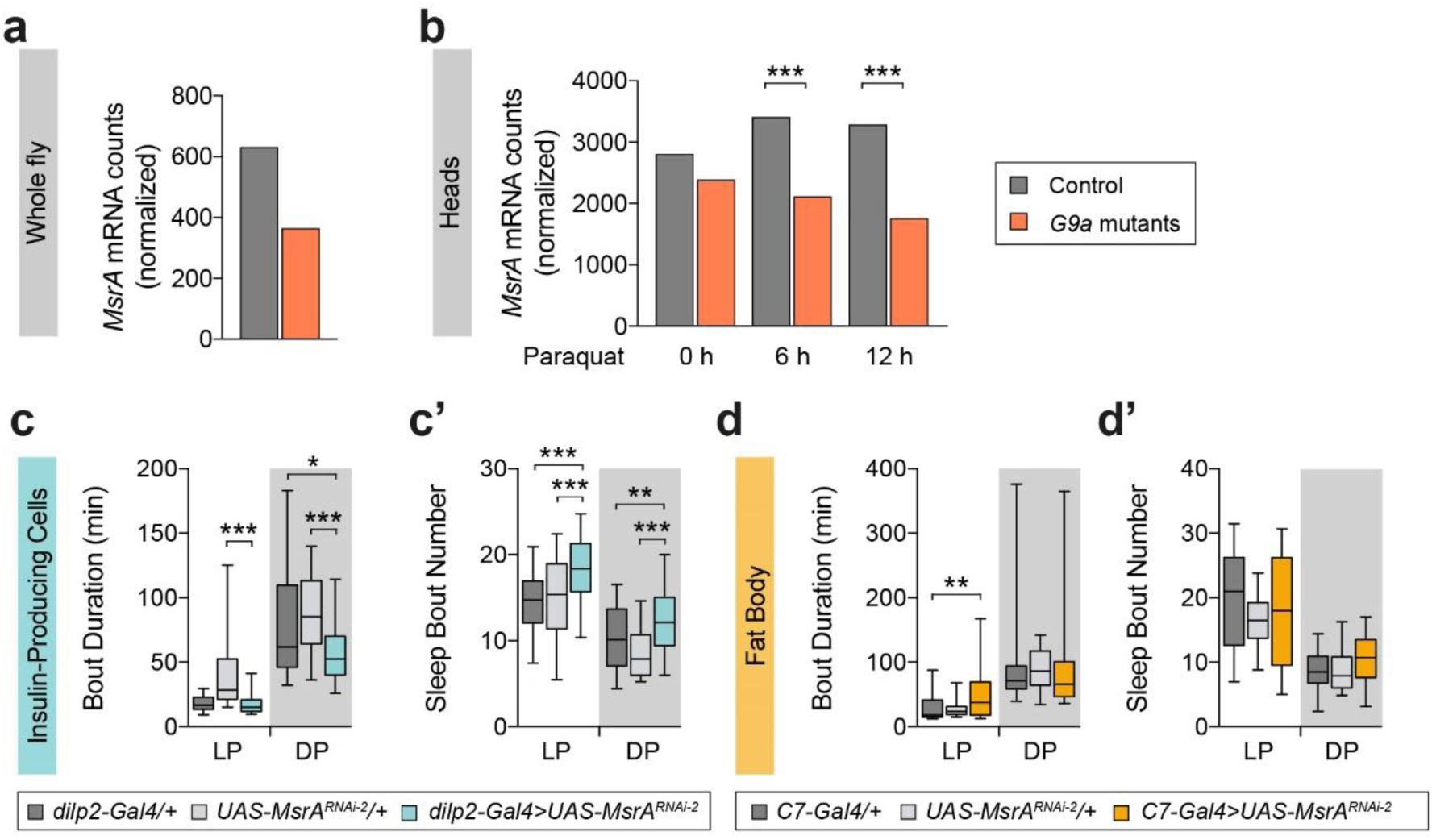
*MsrA* knockdown in insulin-producing cells, but not fat body, leads to fragmented nighttime sleep. **(a)** Normalized RNA-seq counts for *MsrA* in whole *G9a* mutants and isogenic controls^12^. Data represents one sample per genotype and are presented without inferential statistics. **(b)** Normalized RNA-seq counts for *MsrA* in heads isolated from homozygous G9a mutant females 0 h (*p* > 0.05), 6 h (*p* = 4.09e-16), and 12 h (*p* = 1.009e-26) after paraquat treatment relative to isogenic controls^13^. Wald test with Benjamini-Hochberg correction. **(c,d)** Quantification of average duration and **(c’,d’)** number of sleep bouts during the light period (LP, ZT0-12) and dark period (DP, ZT12-24). **(c,c’)** *MsrA* knockdown in IPC neurons (*dilp2-Gal4 > UAS*-*MsrA^RNAi-2^*, n = 70) leads to shorter sleep bouts (*p* = 0.011 and *p* < 0.0001) during the DP, accompanied by an increase in their number (*p* = 0.0093 and *p* < 0.0001) when compared to both parental controls (*dilp2-Gal4/+*, n = 72; *UAS*-*MsrA^RNAi-2^/+*, n = 48). **(d,d’)** *MsrA* knockdown in the fat body (*UAS-Dcr2, C7-Gal4 > UAS*-*MsrA^RNAi-2^*, n = 48) does not cause significant differences in sleep when compared to both parental controls (*UAS-Dcr2, C7-Gal4/+*, n = 48; *UAS*-*MsrA^RNAi-2^/+*, n = 24). Data are represented as boxplots that extend from the 25^th^ to the 75^th^ percentiles with the median indicated. Whiskers indicate the 5^th^ and 95^th^ percentiles. One-way ANOVA with Bonferroni correction or Kruskal-Wallis test with Dunn’s correction according to the distribution of the data. Bonferroni correction for multiple testing. *p*-values are indicated as follows: \**p <* 0.05, \*\**p <* 0.01, \*\*\**p <* 0.001.

**Supplementary Figure 8.**
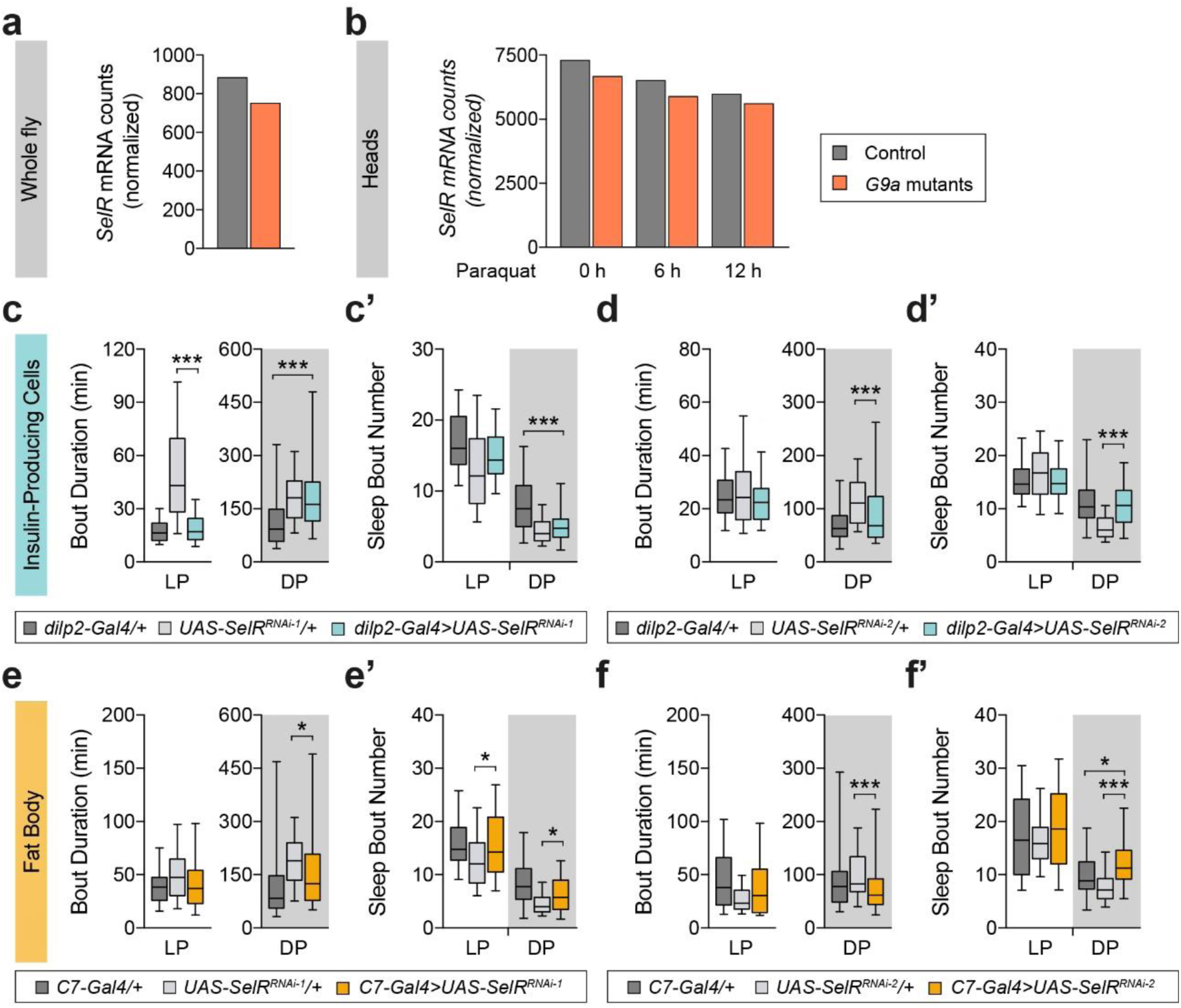
*SelR* (*MsrB*) knockdown in insulin-producing cells or fat body does not phenocopy *G9a*’s nighttime sleep disturbances. **(a)** Normalized RNA-seq counts for *SelR* in whole *G9a* mutants and isogenic controls^12^. Data represents one sample per genotype and are presented without inferential statistics. **(b)** Normalized RNA-seq counts for *SelR* in heads isolated from homozygous G9a mutant females 0 h (*p* > 0.05), 6 h (*p* > 0.05), and 12 h (*p* > 0.05) after paraquat treatment relative to isogenic controls^13^. Wald test with Benjamini-Hochberg correction. **(c-f)** Quantification of average duration and **(c’-f’)** number of sleep bouts during the light period (LP, ZT0-12) and dark period (DP, ZT12-24). **(c,c’)** *SelR* (*MsrB*) knockdown in IPC neurons (*dilp2-Gal4 > UAS*-*SelR^RNAi-1^*, n = 74) shows no significant differences in sleep when compared to both parental controls (*dilp2-Gal4/+*, n = 71; *UAS*-*SelR^RNAi-1^/+*, n = 72). **(d,d’)** *SelR* (*MsrB*) knockdown in IPC neurons (*dilp2-Gal4 > UAS*-SelR*^RNAi-2^*, n = 72) does not lead to significant differences in sleep when compared to both parental controls (*dilp2-Gal4/+*, n = 72; *UAS*-*SelR^RNAi-2^*/+, n = 71). **(e,e’)** *SelR* (*MsrB*) knockdown in the fat body (*UAS-Dcr2, C7-Gal4 > UAS*-*SelR^RNAi-1^*, n = 70) shows no significant differences in sleep when compared to both parental controls (*UAS-Dcr2, C7-Gal4/+*, n = 65; *UAS*-*SelR^RNAi-1^/+*, n = 72). **(f,f’)** *SelR* (*MsrB*) knockdown in the fat body (*UAS-Dcr2, C7-Gal4 > UAS*-*SelR^RNAi-2^*, n = 70) shows an increase in the number of sleep bouts during the DP when compared to both parental controls (*UAS-Dcr2,C7-Gal4/+*, n = 68; *UAS*-*SelR^RNAi-2^/+*, n = 72). Data are represented as boxplots that extend from the 25^th^ to the 75^th^ percentiles with the median indicated. Whiskers indicate the 5^th^ and 95^th^ percentiles. One-way ANOVA with Bonferroni correction or Kruskal-Wallis test with Dunn’s correction according to the distribution of the data. Bonferroni correction for multiple testing. *p*-values are indicated as follows: \**p <* 0.05, \*\**p <* 0.01, \*\*\**p <* 0.001.

**Supplementary Figure 9.**
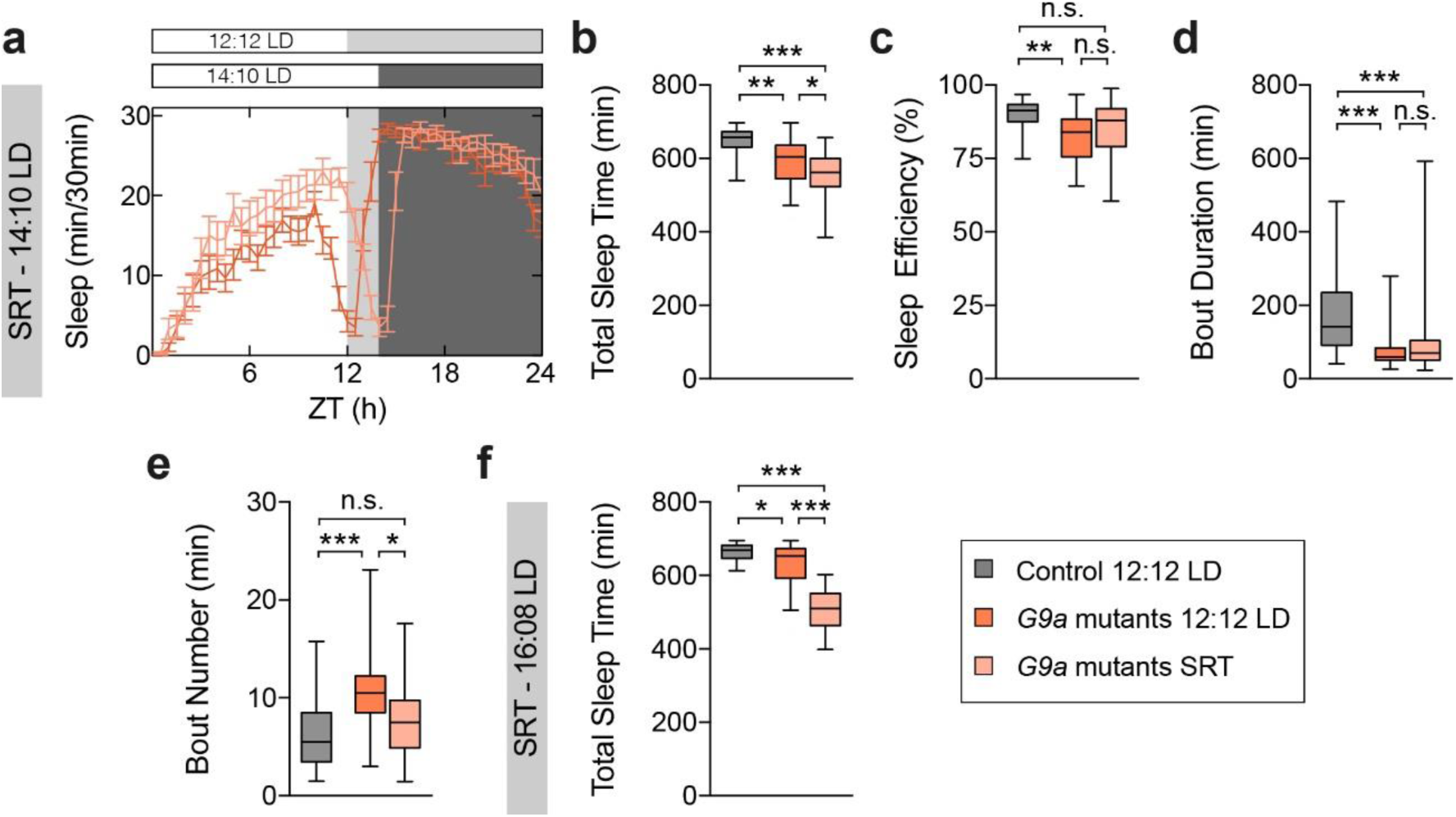
G9a sleep data upon intervention with 14:10 and 16:8 LD Sleep-restriction therapy regimes. **(a)** Representative sleep profiles of *G9a* mutant flies entrained either at 12:12 LD (n = 21) or 14:10 LD (shortened dark period ZT14-24, n = 22) compared to isogenic controls entrained at 12:12 LD. **(b)** Quantification of total sleep time, **(c)** sleep efficiency, **(d)** sleep bout duration, and **(e)** bout number of *G9a* mutant flies entrained either at 12:12 LD (n = 37) or 14:10 LD (n = 38) compared to isogenic controls entrained at 12:12: LD (n = 38) in the dark period (DP). **(f)** Quantification of total sleep time of *G9a* mutant flies entrained at either 12:12 LD (n = 71) or 16:8 LD (n = 69) compared to isogenic controls entrained at 12:12 LD (n = 72). Data are represented as boxplots that extend from the 25^th^ to the 75^th^ percentiles with the median indicated. Whiskers indicate the 5^th^ and 95^th^ percentiles. Kruskal-Wallis test with Dunn’s correction. Bonferroni correction for multiple testing. *p*-values are indicated as follows: \**p <* 0.05, \*\**p <* 0.01, \*\*\**p <* 0.001.

## References

1. Bruni, O., Breda, M., Mammarella, V., Mogavero, M.P. & Ferri, R. Sleep and circadian disturbances in children with neurodevelopmental disorders. Nat Rev Neurol 21, 103– 120 (2025).

2. Kleefstra, T., Brunner, H.G., Amiel, J., Oudakker, A.R., Nillesen, W.M., Magee, A., Geneviève, D., Cormier-Daire, V., van Esch, H., Fryns, J.P., Hamel, B.C., Sistermans, E.A., de Vries, B.B. & van Bokhoven, H. Loss-of-function mutations in euchromatin histone methyl transferase 1 (EHMT1) cause the 9q34 subtelomeric deletion syndrome. Am J Hum Genet 79, 370–377 (2006).

3. Kleefstra, T., Smidt, M., Banning, M.J., Oudakker, A.R., Van Esch, H., de Brouwer, A.P., Nillesen, W., Sistermans, E.A., Hamel, B.C., de Bruijn, D., Fryns, J.P., Yntema, H.G., Brunner, H.G., de Vries, B.B. & van Bokhoven, H. Disruption of the gene Euchromatin Histone Methyl Transferase1 (Eu-HMTase1) is associated with the 9q34 subtelomeric deletion syndrome. J Med Genet 42, 299–306 (2005).

4. Bouman, A., Gaasterland, C.M.W., Sloof-Enthoven, C., Draksler, T.Z., Rots, D., Vermeulen-Kalk, K., Geelen, J.M., Morison, L.D., Morgan, A.T., Wicher, D., Rivero, S., Fernandez-Ulibarri, I., Drake, J., O’Donnell Luria, A., Pickup, L., Shalhoub, C., Milani, D., Hennekam, R.C., Tumiene, B., Dies, K.A., Garavelli, L., Bedeschi, M.F., Danieli, A., van Renssen, L.V., Palmer, E.E., Grosdemouge, I., Hadzsiev, K., Ousager, L.B., Frazier, Z., Chopra, M., Szakszon, K., Ewans, L., Srivastava, S., Balbo, N., Caterino, E., Schenck, A., Smith, R., Boonstra, F.N., van Till, S.A.L., Vasireddi, S.K., Brian Chung, H.Y., Klein Haneveld, M.J., Vyshka, K., Group, E.R.N.I.G.W. & Kleefstra, T. International clinical evidence-based guideline for Kleefstra syndrome. Genet Med 28, 102070 (2026).

5. Bouman, A., Geelen, J.M., Kummeling, J., Schenck, A., van der Zwan, Y.G., Klein, W.M. & Kleefstra, T. Growth, body composition, and endocrine-metabolic profiles of individuals with Kleefstra syndrome provide directions for clinical management and translational studies. Am J Med Genet A 194, e63472 (2024).

6. Kleefstra, T., Kramer, J.M., Neveling, K., Willemsen, M.H., Koemans, T.S., Vissers, L.E., Wissink-Lindhout, W., Fenckova, M., van den Akker, W.M., Kasri, N.N., Nillesen, W.M., Prescott, T., Clark, R.D., Devriendt, K., van Reeuwijk, J., de Brouwer, A.P., Gilissen, C., Zhou, H., Brunner, H.G., Veltman, J.A., Schenck, A. & van Bokhoven, H. Disruption of an EHMT1-associated chromatin-modification module causes intellectual disability. Am J Hum Genet 91, 73–82 (2012).

7. Rots, D., Bouman, A., Yamada, A., Levy, M., Dingemans, A.J.M., de Vries, B.B.A., Ruiterkamp-Versteeg, M., de Leeuw, N., Ockeloen, C.W., Pfundt, R., de Boer, E., Kummeling, J., van Bon, B., van Bokhoven, H., Kasri, N.N., Venselaar, H., Alders, M., Kerkhof, J., McConkey, H., Kuechler, A., Elffers, B., van Beeck Calkoen, R., Hofman, S., Smith, A., Valenzuela, M.I., Srivastava, S., Frazier, Z., Maystadt, I., Piscopo, C., Merla, G., Balasubramanian, M., Santen, G.W.E., Metcalfe, K., Park, S.M., Pasquier, L., Banka, S., Donnai, D., Weisberg, D., Strobl-Wildemann, G., Wagemans, A., Vreeburg, M., Baralle, D., Foulds, N., Scurr, I., Brunetti-Pierri, N., van Hagen, J.M., Bijlsma, E.K., Hakonen, A.H., Courage, C., Genevieve, D., Pinson, L., Forzano, F., Deshpande, C., Kluskens, M.L., Welling, L., Plomp, A.S., Vanhoutte, E.K., Kalsner, L., Hol, J.A., Putoux, A., Lazier, J., Vasudevan, P., Ames, E., O’Shea, J., Lederer, D., Fleischer, J., O’Connor, M., Pauly, M., Vasileiou, G., Reis, A., Kiraly-Borri, C., Bouman, A., Barnett, C., Nezarati, M., Borch, L., Beunders, G., Ozcan, K., Miot, S., Volker-Touw, C.M.L., van Gassen, K.L.I., Cappuccio, G., Janssens, K., Mor, N., Shomer, I., Dominissini, D., Tedder, M.L., Muir, A.M., Sadikovic, B., Brunner, H.G., Vissers, L., Shinkai, Y. & Kleefstra, T. Comprehensive EHMT1 variants analysis broadens genotype-phenotype associations and molecular mechanisms in Kleefstra syndrome. Am J Hum Genet 111, 1605–1625 (2024).

8. Schmidt, S., Nag, H.E., Hunn, B.S., Houge, G. & Hoxmark, L.B. A structured assessment of motor function and behavior in patients with Kleefstra syndrome. Eur J Med Genet 59, 240–248 (2016).

9. Vermeulen, K., de Boer, A., Janzing, J.G.E., Koolen, D.A., Ockeloen, C.W., Willemsen, M.H., Verhoef, F.M., van Deurzen, P.A.M., van Dongen, L., van Bokhoven, H., Egger, J.I.M., Staal, W.G. & Kleefstra, T. Adaptive and maladaptive functioning in Kleefstra syndrome compared to other rare genetic disorders with intellectual disabilities. Am J Med Genet A 173, 1821–1830 (2017).

10. Morison, L.D., Kennis, M.G.P., Rots, D., Bouman, A., Kummeling, J., Palmer, E., Vogel, A.P., Liegeois, F., Brignell, A., Srivastava, S., Frazier, Z., Milnes, D., Goel, H., Amor, D.J., Scheffer, I.E., Kleefstra, T. & Morgan, A.T. Expanding the phenotype of Kleefstra syndrome: speech, language and cognition in 103 individuals. J Med Genet 61, 578–585 (2024).

11. Kramer, J.M., Kochinke, K., Oortveld, M.A., Marks, H., Kramer, D., de Jong, E.K., Asztalos, Z., Westwood, J.T., Stunnenberg, H.G., Sokolowski, M.B., Keleman, K., Zhou, H., van Bokhoven, H. & Schenck, A. Epigenetic regulation of learning and memory by Drosophila EHMT/G9a. PLoS biology 9, e1000569 (2011).

12. Merkling, S.H., Bronkhorst, A.W., Kramer, J.M., Overheul, G.J., Schenck, A. & Van Rij, R.P. The epigenetic regulator G9a mediates tolerance to RNA virus infection in Drosophila. PLoS pathogens 11, e1004692 (2015).

13. Riahi, H., Brekelmans, C., Foriel, S., Merkling, S.H., Lyons, T.A., Itskov, P.M., Kleefstra, T., Ribeiro, C., van Rij, R.P., Kramer, J.M. & Schenck, A. The histone methyltransferase G9a regulates tolerance to oxidative stress-induced energy consumption. PLoS biology 17, e2006146 (2019).

14. Riahi, H., Fenckova, M., Goruk, K.J., Schenck, A. & Kramer, J.M. The epigenetic regulator G9a attenuates stress-induced resistance and metabolic transcriptional programs across different stressors and species. BMC Biol 19, 112 (2021).

15. Coll-Tane, M., Eidhof, I., Han, J., Raun, N., van Renssen, L.V., Fisher, S.E., Kayser, M.S., Kleefstra, T., Pillen, S., Hudac, C.M., Mayneris-Perxachs, J., Klein, M., Koene, S., Castells-Nobau, A. & Schenck, A. Conserved sleep disturbances in FOXP1 syndrome originate from developmental dysregulation of peptidergic signaling. J Clin Invest 136(2026).

16. Coll-Tane, M., Gong, N.N., Belfer, S.J., van Renssen, L.V., Kurtz-Nelson, E.C., Szuperak, M., Eidhof, I., van Reijmersdal, B., Terwindt, I., Durkin, J., Verheij, M.M.M., Kim, C.N., Hudac, C.M., Nowakowski, T.J., Bernier, R.A., Pillen, S., Earl, R.K., Eichler, E.E., Kleefstra, T., Kayser, M.S. & Schenck, A. The CHD8/CHD7/Kismet family links blood-brain barrier glia and serotonin to ASD-associated sleep defects. Sci Adv 7(2021).

17. Coll-Tané, M., Krebbers, A., Castells-Nobau, A., Zweier, C. & Schenck, A. Intellectual disability and autism spectrum disorders ‘on the fly’: insights from Drosophila. Dis Model Mech 12(2019).

18. Chakravarti, L., Moscato, E.H. & Kayser, M.S. Unraveling the Neurobiology of Sleep and Sleep Disorders Using Drosophila. Curr Top Dev Biol 121, 253–285 (2017).

19. Simonds, J.F. & Parraga, H. Prevalence of sleep disorders and sleep behaviors in children and adolescents. J Am Acad Child Psychiatry 21, 383–388 (1982).

20. Simonds, J.F. & Parraga, H. Sleep behaviors and disorders in children and adolescents evaluated at psychiatric clinics. J Dev Behav Pediatr 5, 6–10 (1984).

21. Wiggs, L. & Stores, G. Behavioural treatment for sleep problems in children with severe learning disabilities and challenging daytime behaviour: effect on sleep patterns of mother and child. J Sleep Res 7, 119–126 (1998).

22. Medicine, A.A.o.S. International classification of sleep disorders—third edition (ICSD-3). AASM Resour Libr 281, 2313 (2014).

23. Watanabe, K., Stringer, S., Frei, O., Umićević Mirkov, M., de Leeuw, C., Polderman, T.J.C., van der Sluis, S., Andreassen, O.A., Neale, B.M. & Posthuma, D. A global overview of pleiotropy and genetic architecture in complex traits. Nat Genet 51, 1339–1348 (2019).

28. Dietzl, G., Chen, D., Schnorrer, F., Su, K.C., Barinova, Y., Fellner, M., Gasser, B., Kinsey, K., Oppel, S., Scheiblauer, S., Couto, A., Marra, V., Keleman, K. & Dickson, B.J. A genome-wide transgenic RNAi library for conditional gene inactivation in Drosophila. Nature 448, 151–156 (2007).

29. Joiner, W.J., Crocker, A., White, B.H. & Sehgal, A. Sleep in Drosophila is regulated by adult mushroom bodies. Nature 441, 757–760 (2006).

30. Chen, K.F., Lowe, S., Lamaze, A., Krätschmer, P. & Jepson, J. Neurocalcin regulates nighttime sleep and arousal in Drosophila. Elife 8(2019).

31. Jenett, A., Rubin, G.M., Ngo, T.T., Shepherd, D., Murphy, C., Dionne, H., Pfeiffer, B.D., Cavallaro, A., Hall, D., Jeter, J., Iyer, N., Fetter, D., Hausenfluck, J.H., Peng, H., Trautman, E.T., Svirskas, R.R., Myers, E.W., Iwinski, Z.R., Aso, Y., DePasquale, G.M., Enos, A., Hulamm, P., Lam, S.C., Li, H.H., Laverty, T.R., Long, F., Qu, L., Murphy, S.D., Rokicki, K., Safford, T., Shaw, K., Simpson, J.H., Sowell, A., Tae, S., Yu, Y. & Zugates, C.T. A GAL4-driver line resource for Drosophila neurobiology. Cell Rep 2, 991–1001 (2012).

32. Hill, V.M., O’Connor, R.M. & Shirasu-Hiza, M. Tired and stressed: Examining the need for sleep. Eur J Neurosci 51, 494–508 (2020).

33. Brown, E.B., Shah, K.D., Faville, R., Kottler, B. & Keene, A.C. Drosophila insulin-like peptide 2 mediates dietary regulation of sleep intensity. PLoS Genet 16, e1008270 (2020).

34. Crocker, A., Shahidullah, M., Levitan, I.B. & Sehgal, A. Identification of a neural circuit that underlies the effects of octopamine on sleep:wake behavior. Neuron 65, 670–681 (2010).

35. Rajan, A. & Perrimon, N. Drosophila cytokine unpaired 2 regulates physiological homeostasis by remotely controlling insulin secretion. Cell 151, 123–137 (2012).

36. Zhang, Y. & Xi, Y. Fat body development and its function in energy storage and nutrient sensing in Drosophila melanogaster. Journal of Tissue Science & Engineering 6, 1 (2015).

37. McGuire, S.E., Mao, Z. & Davis, R.L. Spatiotemporal gene expression targeting with the TARGET and gene-switch systems in Drosophila. Sci STKE 2004, pl6 (2004).

38. Stadtman, E.R. Oxidation of free amino acids and amino acid residues in proteins by radiolysis and by metal-catalyzed reactions. Annu Rev Biochem 62, 797–821 (1993).

39. Vogt, W. Oxidation of methionyl residues in proteins: tools, targets, and reversal. Free Radic Biol Med 18, 93–105 (1995).

40. Chao, C.C., Ma, Y.S. & Stadtman, E.R. Modification of protein surface hydrophobicity and methionine oxidation by oxidative systems. Proc Natl Acad Sci U S A 94, 2969–2974 (1997).

41. Lee, B.C. & Gladyshev, V.N. The biological significance of methionine sulfoxide stereochemistry. Free Radic Biol Med 50, 221–227 (2011).

42. Rebrin, I. & Sohal, R.S. Pro-oxidant shift in glutathione redox state during aging. Adv Drug Deliv Rev 60, 1545–1552 (2008).

43. Hill, V.M., O’Connor, R.M., Sissoko, G.B., Irobunda, I.S., Leong, S., Canman, J.C., Stavropoulos, N. & Shirasu-Hiza, M. A bidirectional relationship between sleep and oxidative stress in Drosophila. PLoS biology 16, e2005206 (2018).

44. Albrecht, S.C., Barata, A.G., Grosshans, J., Teleman, A.A. & Dick, T.P. In vivo mapping of hydrogen peroxide and oxidized glutathione reveals chemical and regional specificity of redox homeostasis. Cell Metab 14, 819–829 (2011).

45. Terzi, A., Ngo, K.J. & Mourrain, P. Phylogenetic conservation of the interdependent homeostatic relationship of sleep regulation and redox metabolism. Journal of Comparative Physiology B 194, 241–252 (2024).

46. Tian, Y., Liu, X. & Liu, D. Physiological roles of reactive oxygen species in sleep homeostasis. Free Radic Biol Med 242, 220–236 (2026).

47. Liu, L., Zhang, K., Sandoval, H., Yamamoto, S., Jaiswal, M., Sanz, E., Li, Z., Hui, J., Graham, B.H., Quintana, A. & Bellen, H.J. Glial lipid droplets and ROS induced by mitochondrial defects promote neurodegeneration. Cell 160, 177–190 (2015).

48. Bailey, A.P., Koster, G., Guillermier, C., Hirst, E.M., MacRae, J.I., Lechene, C.P., Postle, A.D. & Gould, A.P. Antioxidant Role for Lipid Droplets in a Stem Cell Niche of Drosophila. Cell 163, 340–353 (2015).

49. Damri, O., Natour, S., Asslih, S. & Agam, G. Does treatment with autophagy-enhancers and/or ROS-scavengers alleviate behavioral and neurochemical consequences of low-dose rotenone-induced mild mitochondrial dysfunction in mice? Mol Psychiatry (2023).

50. Shaposhnikov, M.V., Zemskaya, N.V., Koval, L.A., Schegoleva, E.V., Zhavoronkov, A. & Moskalev, A.A. Effects of N-acetyl-L-cysteine on lifespan, locomotor activity and stress-resistance of 3 Drosophila species with different lifespans. Aging (Albany NY) 10, 2428– 2458 (2018).

51. Kumar, R.A., Koc, A., Cerny, R.L. & Gladyshev, V.N. Reaction mechanism, evolutionary analysis, and role of zinc in Drosophila methionine-R-sulfoxide reductase. J Biol Chem 277, 37527–37535 (2002).

52. Miller, C.B., Espie, C.A., Epstein, D.R., Friedman, L., Morin, C.M., Pigeon, W.R., Spielman, A.J. & Kyle, S.D. The evidence base of sleep restriction therapy for treating insomnia disorder. Sleep Med Rev 18, 415–424 (2014).

53. Belfer, S.J., Bashaw, A.G., Perlis, M.L. & Kayser, M.S. A Drosophila model of sleep restriction therapy for insomnia. Mol Psychiatry (2019).

54. Zdolsek Draksler, T., Bouman, A., Gucek, A., Novak, E., Burger, P., Colin, F. & Kleefstra, T. Exploring Kleefstra syndrome cohort phenotype characteristics: Prevalence insights from caregiver-reported outcomes. Eur J Med Genet 72, 104974 (2024).

55. Frazier, Z.J., Kilic, S., Osika, H., Mo, A., Quinn, M., Ballal, S., Katz, T., Shearer, A.E., Horlbeck, M.A., Pais, L.S., Dies, K.A., O’Donnell-Luria, A., Kossowsky, J., Lipton, J.O., Kleefstra, T. & Srivastava, S. Novel Phenotypes and Genotype-Phenotype Correlations in a Large Clinical Cohort of Patients With Kleefstra Syndrome. Clin Genet 107, 636–645 (2025).

56. Haseley, A., Wallis, K. & DeBrosse, S. Kleefstra syndrome: Impact on parents. Disabil Health J 14, 101018 (2021).

57. Rennick-Zuefle, K., Weljie, A.M. & Kain, P. The interplay of insulin signaling and neurotransmitters on sleep in Drosophila. Front Neuroendocrinol 81, 101252 (2026).

58. Jones, S.G., Bouman, A., Raun, N., van Genugten, E.A.J., Martínez-Blázquez, I., Kampshoff, F., Doorduin, J., Geelen, J., Bruining, H., Vermeulen-Kalk, K., Miot, S., Geneviève, D., Aarntzen, E.H.J.G., Coll-Tané, M., Kleefstra, T. & Schenck, A. Metabolic collapse as a mechanism of developmental regression: convergent evidence from Kleefstra syndrome [18F]FDG-PET/CT imaging and Drosophila modelling bioRxiv (2026).

59. Dissel, S., Angadi, V., Kirszenblat, L., Suzuki, Y., Donlea, J., Klose, M., Koch, Z., English, D., Winsky-Sommerer, R., van Swinderen, B. & Shaw, P.J. Sleep restores behavioral plasticity to Drosophila mutants. Curr Biol 25, 1270–1281 (2015).

60. Krugliakova, E., Breuer, F., Adelhofer, N., Alonso, A., Besedovsky, L., Murphy, K., Peters, E., Raczek, K., Rasch, B., Salvesen, L., Snipes, S., Schoch, S., Schreiner, T., Wassing, R., Bergmann, T.O. & Dresler, M. Hacking the functions of sleep: noninvasive approaches to stimulate sleep neurophysiology. Physiol Rev 106, 675–749 (2026).

61. Sharon, O., Ben Simon, E., Shah, V.D., Desel, T. & Walker, M.P. The new science of sleep: From cells to large-scale societies. PLoS biology 22, e3002684 (2024).

62. Paruthi, S., Brooks, L.J., D’Ambrosio, C., Hall, W.A., Kotagal, S., Lloyd, R.M., Malow, B.A., Maski, K., Nichols, C., Quan, S.F., Rosen, C.L., Troester, M.M. & Wise, M.S. Recommended Amount of Sleep for Pediatric Populations: A Consensus Statement of the American Academy of Sleep Medicine. J Clin Sleep Med 12, 785–786 (2016).

63. Mollayeva, T., Thurairajah, P., Burton, K., Mollayeva, S., Shapiro, C.M. & Colantonio, A. The Pittsburgh sleep quality index as a screening tool for sleep dysfunction in clinical and non-clinical samples: A systematic review and meta-analysis. Sleep Med Rev 25, 52– 73 (2016).

64. de Leeuw, C.A., Mooij, J.M., Heskes, T. & Posthuma, D. MAGMA: generalized gene-set analysis of GWAS data. PLoS Comput Biol 11, e1004219 (2015).

65. Rynes, J., Donohoe, C.D., Frommolt, P., Brodesser, S., Jindra, M. & Uhlirova, M. Activating transcription factor 3 regulates immune and metabolic homeostasis. Mol Cell Biol 32, 3949–3962 (2012).

66. Shaw, P.J., Cirelli, C., Greenspan, R.J. & Tononi, G. Correlates of sleep and waking in Drosophila melanogaster. Science (New York, N.Y.) 287, 1834–1837 (2000).

67. Hendricks, J.C., Finn, S.M., Panckeri, K.A., Chavkin, J., Williams, J.A., Sehgal, A. & Pack, A.I. Rest in Drosophila is a sleep-like state. Neuron 25, 129–138 (2000).

68. Donelson, N.C., Kim, E.Z., Slawson, J.B., Vecsey, C.G., Huber, R. & Griffith, L.C. High-resolution positional tracking for long-term analysis of Drosophila sleep and locomotion using the "tracker" program. PLoS One 7, e37250 (2012).

69. Ritchie, M.E., Phipson, B., Wu, D., Hu, Y., Law, C.W., Shi, W. & Smyth, G.K. limma powers differential expression analyses for RNA-sequencing and microarray studies. Nucleic Acids Research 43, e47–e47 (2015).

70. Haynes, P.R., Pyfrom, E.S., Li, Y., Stein, C., Cuddapah, V.A., Jacobs, J.A., Yue, Z. & Sehgal, A. A neuron-glia lipid metabolic cycle couples daily sleep to mitochondrial homeostasis. Nat Neurosci 27, 666–678 (2024).

