## Supplemental Table 1 for "The evolutionarily conserved EHMT1/G9a histone methyltransferase family regulates sleep maintenance through ROS homeostasis in insulin-producing cells"

| **Trait** | *p*-value | Domain | Atlas ID | PMID | Year | N |
| --- | --- | --- | --- | --- | --- | --- |
| Chronotype | **0.004522** | Psychiatric | 1141 | 27494321 | 2016 | 128266 |
| Daytime sleepiness / dozing | 0.034 | Psychiatric | 3790 | 30804565 | 2019 | 385333 |
| Excessive daytime sleepiness | **0.00303** | Psychiatric | 1175 | 27992416 | 2017 | 111648 |
| Extreme chronotype | 0.078 | Psychiatric | 1174 | 26955885 | 2016 | 100420 |
| Frequent insomnia symptoms | **0.00406** | Neurological | 4290 | 30804566 | 2019 | 237627 |
| Insomnia | 0.0066 | Neurological | 3786 | 30804565 | 2019 | 386533 |
| Long sleep | 0.503 | Psychiatric | 4273 | 30846698 | 2019 | 339926 |
| Number of sleep episodes | 0.021 | Psychiatric | 4252 | 30952852 | 2019 | 84810 |
| Short sleep | **0.0016** | Psychiatric | 4274 | 30846698 | 2019 | 411934 |
| Sleep duration | **1.97E-06** | Psychiatric | 4272 | 30846698 | 2019 | 446118 |
| Sleep efficiency | **0.0028** | Psychiatric | 4255 | 30952852 | 2019 | 84810 |

**Supplementary Table 1 – Common variation in *EHMT1* is significantly associated with multiple sleep traits in the general population.** Common variants contained in *EHMT1* are significantly associated with the sleep traits “chronotype”, “excessive daytime sleepiness”, “sleep efficiency”, “sleep duration”, “short sleep”, and “frequent insomnia symptoms”. Bonferroni correction for multiple comparisons was applied to the significance level (*p* < 0.0045). Values in bold withstand correction for multiple testing.
